# The high-energy transition state of a membrane transporter

**DOI:** 10.1101/2020.04.17.047373

**Authors:** Gerard H. M. Huysmans, Didar Ciftci, Xiaoyu Wang, Scott C. Blanchard, Olga Boudker

**Author notes:** correspondence: Gerard Huysmans,; Olga Boudker.

## Abstract

Membrane transporters mediate cellular uptake of nutrients, signaling molecules and drugs. Their overall mechanisms are often well understood, but the structural features setting their rates are mostly unknown. Earlier single-molecule fluorescence imaging of a model glutamate transporter homologue suggested that the slow conformational transition from the outward- to the inward-facing state, when the bound substrate is translocated from the extracellular to the cytoplasmic side of the membrane, is rate-limiting to transport. Here, we aim to gain insight into the structure of the high-energy transition state that limits the rate of this critical isomerization reaction. Using bioinformatics, we identify gain-of-function mutants of the transporter and apply linear free energy relationship analysis to infer that the transition state structurally resembles the inward-facing conformation. Based on these analyses, we propose an approach for allosteric modulation of these transporters.

---

The fluxes of small molecules across biological membranes mediated by membrane-embedded transporters are essential to life, and their dysregulation leads to numerous diseases ^1^. High-resolution structures of transporters have revealed conformations that expose substrate-binding sites to the opposite sides of the membrane, providing a structural rationale for the alternating access mechanism ^2^. In contrast, little is known about the structures of high-energy transition states (TSs) that are essential to understand the kinetic mechanisms and to develop drugs that modulate transporters activity ^3^. While many computational approaches are used to map the energy landscapes of the transporters ^4-16^, experimental data reporting on TSs are scarce ^17,18^.

TSs are only transiently populated and cannot be studied by direct structural methods However, linear free energy relationship (LFER) analyses have been developed to characterize TSs of specific enzyme reactions ^19-21^, protein folding ^22-26^ and channel opening ^27-29^. LFERs correlate the effects of amino acid substitutions and ligands on the reaction rates to the changes in the equilibrium constant for an observed transition. From these correlations, one can infer whether the TS is structurally more similar to the initial or the final conformation ^30-32^. Here, we use LFERs to characterize the structure of the TS during substrate translocation by Glt_Ph_, an aspartate/sodium symporter from the hyperthermophilic archaebacterium *Pyrococcus horikoshii*.

Glt_Ph_ is an extensively studied homologue of human excitatory amino acid transporters (EAATs) ^33^. It utilizes the physiological trans-membrane sodium (Na^+^) gradient by symporting one aspartate (l-Asp) and three Na^+^ ions ^34-36^. From a structural perspective, the mechanism of Glt_Ph_ is well understood. Glt_Ph_ is a homotrimer, with each protomer consisting of a trimerization scaffold domain and a transport domain, containing the l-Asp- and Na^+^-binding sites. The solutes are delivered across the bilayer upon a ∼15 Å “elevator” movement of the transport domain along the membrane normal from an outward-facing state (OFS) to an inward-facing state (IFS) (**Figure 1A**) ^37-47^. During this transition, the transport domain forms two alternative interfaces with the scaffold involving pseudo-symmetric helical hairpins 1 and 2 (HP1 and 2) ^46,48,49^. In the OFS, HP1 is sandwiched between the transport domain and the scaffold, while HP2 lines the extracellular surface of the protomer. In the IFS, HP2 is positioned on the interface, and HP1 faces the cytosol (**Figure 1A**). HP2 forms a lid over the substrate-binding site and serves as the extracellular gate ^34,50-55^. The release of l-Asp and Na^+^-ions in the IFS occurs via an incompletely understood mechanism and leads to the collapse of HP2 onto the substrate-binding site ^56,57^. The cycle is completed when the transport domain returns to the OFS (**Figure 1B**). Time-resolved single-molecule Forster resonance energy transfer (smFRET) recordings have been instrumental in establishing the key features of the transport domains movements ^37,38,41^. These studies suggested that the translocation of the substrate-loaded transport domain from the OFS to the IFS is the rate-limited step of the cycle, but the nature of the high-energy barrier that determines the translocation rate remained unknown.

**Figure 1:**
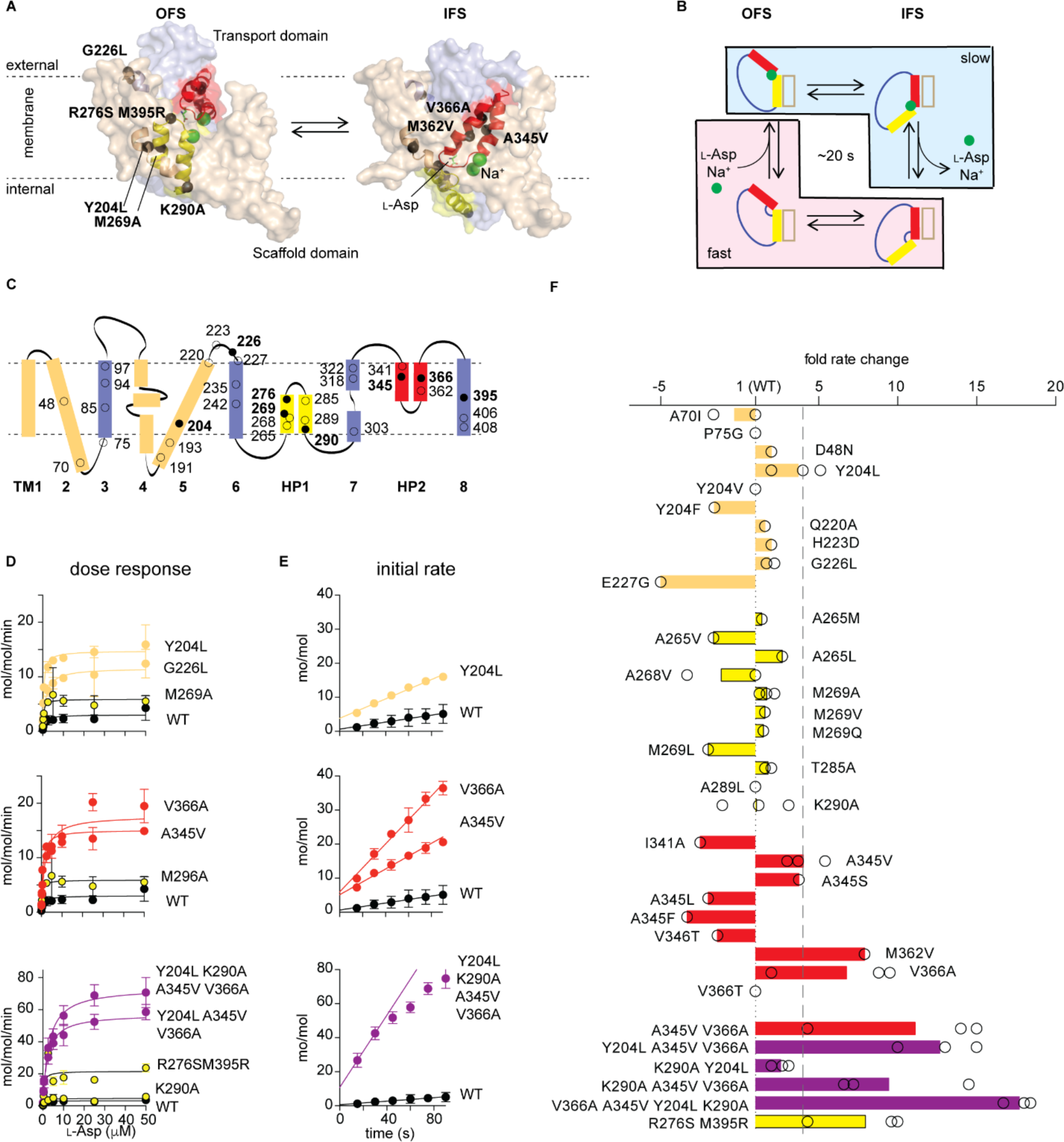
Gain-of-function Glt_Ph_ variants. (**A**) A Glt_Ph_ protomer in the OFS and the IFS is shown in semi-transparent surface representation with the scaffold and transport domains colored wheat and blue, respectively. HP1 (yellow) and HP2 (red) are emphasized as cartoons. Bound l-Asp and Na^+^ ions are shown as green sticks and spheres, respectively. Select amino acids mutated in this study are shown as black spheres. Color coding is the same in all panels. (**B**) Schematic representation of the Glt_Ph_ transport cycle showing one protomer. Comparatively rapid and slow steps of the cycle are shaded pink and blue, respectively. (**C**) Topology of a Glt_Ph_ protomer showing tested mutation sites (circles) with filled circles corresponding to those shown in A. (**D, E**) Representative examples of dose-response curves and time courses. Lines through the dose-response curves are fits to the Michaelis-Menten equation; time courses are fitted to linear equations. Shown are means and standard errors over at least three independent repeats. (**F**) Changes in the initial uptake rates of the variants relative to WT Glt_Ph_. Circles are averages of technical triplicates. Bars are means over independent repeats. The dashed line marks a four-fold increase of the initial rate. In D to F, data are colored according to the structural elements where the mutations are located with combination mutants colored purple. *See also Supplementary Figure 1 and Supplementary Tables 1 and 2.*

Here, we generated a repertoire of gain-of-function Glt_Ph_ mutants, in part, through the analysis of systematic amino acid sequence variations among glutamate transporters from organisms living at different temperatures. Our fastest mutant showed the substrate uptake rate ∼20-fold higher than the wild type (WT) transporter. Close examination of a representative set of these mutants corroborated previous findings that the increased rate of the OFS to the IFS transitions of the substrate-loaded transport domain was necessary to increase uptake rate (Akyuz et al., 2015), but also showed that the more dynamic mutants benefited from reduced substrate affinity. At the single-molecule level, the transport domain dynamics of the WT Glt_Ph_ and more dynamic mutants were highly heterogeneous showing a broad distribution of transition frequencies between individual molecules in the population. These apparent dynamic modes parallel the activity modes observed in a recently developed single-molecule transport assay (Ciftci et al., 2020, Science Advances, in review after revisions).

We focused on the dynamic mode in which WT Glt_Ph_ and examined gain-of-function mutants spent most of their time to probe the transition-state structure. In this predominant mode, all of the mutations that increased transport domain dynamics increased the rate of the OFS to the IFS transitions, but most had no effect on the rate of the reverse reaction. Based on these observations, the LFER analysis predicts that the high-energy TS structurally resembles the IFS. Thus, our data suggest that the transport domain might make multiple attempts at reaching the IFS-like TS during the OFS residence before progressing to the stable observable IFS. We further propose that small molecules with a higher affinity for the IFS than the OFS would also have a higher affinity for the TS. These molecules would, therefore, lower the height of the energy barrier of the OFS to the IFS transition and speed up transport. Such positive allosteric modulators of human transporter may offer hitherto unexplored therapeutic avenues.

## Results

### Gain-of-function mutations

Originating from a hyperthermophile, Glt_Ph_ is a slow transporter ^34,36^ (**Figure 1B**). This property is in line with observations that enzymes from thermophilic organisms, which evolved to be stable and functional at high temperatures, show low activity at ambient temperature ^58,59^. Consistently, glutamate transporters from mesophilic bacteria are more active than Glt_Ph_ ^60-63^. Also, structurally very similar human EAATs ^40^, are ∼20 to 10000-fold faster ^33^. Interestingly, a “humanizing” R276S/M395R mutation in Glt_Ph_, which moves an arginine proximal to the substrate binding site from its location in Glt_Ph_ to that in EAATs, confers an increased transport rate ^64^. Inspired by these considerations, we searched for additional gain-of-function mutations by identifying potential evolutionary adaptations to achieve higher activity at lower temperatures. We constructed a multiple sequence alignment of prokaryotic glutamate transporters sorted by the optimum growth temperature of their species of origin and looked for systematic variations between sequences from hyperthermophiles, thermophiles, mesophiles, and psychrophiles (**Supplementary Figure 1**). Overall, we did not observe highly significant systematic changes across all sequences, suggesting that different mutations occurred in different evolutionary lineages. Nevertheless, when we mapped residues with positive global differences scores (**Methods and Supplementary Table 1**) onto the structure of Glt_Ph_, we found that the majority formed two clusters in the transport domain centered on HP1 and HP2 (**Supplementary Figure 1 and Supplementary Table 1**). Among these, sites with a preference for smaller amino acids in hyperthermophiles were more likely to be taken by larger amino acids in mesophiles and psychrophiles and *vice versa* (**Supplementary Table 1**). These observations suggest that packing interactions within the transport domain might play a role during temperature adaptation, consistent with earlier studies where thermophilic and psychrophilic enzymes were, respectively, more rigid and more dynamic than their mesophilic counterparts ^65,66^.

Using this analysis, but also expanding into other regions of the protein (**Supplementary Figure 1**), we selected 30 sites (**Figure 1C**) and tested the transport activity of 44 single mutants reconstituted into proteoliposomes (**Supplementary Table 2**). Most, but not all, showed *K*_*M*_ values similar to the WT *K*_*M*_ of 0.5 ± 0.04 μM, but different maximal rates (**Figure 1D, Supplementary Table 2**). We subsequently measured the initial transport rates of all mutants at l-Asp concentrations of 5-fold over the *K*_*M*_ values (**Figures 1D, F**). Five mutations increased the transport rate at least four-fold (**Figure 1F, Supplementary Table 2**). Four of these (A345V, A345S, M362A, and V366A) were in HP2 with A345 and V366 sites identified by the bioinformatics analysis (**Supplementary Figure 1, Supplementary Table 1**). All four increased *K*_*M*_ at least two-fold. The fifth mutation, Y204L Glt_Ph_, located to the kink of TM5 in the scaffold domain (**Figure 1A, F, Supplementary Table 2**). This site did not receive a high score in our analysis because ∼80% of the bacterial sequences already have an aliphatic residue at this position, but is notable because its flexibility might facilitate transport domain movements ^67^. Mutations elsewhere failed to boost transport (**Figure 1F, Supplementary Table 2**). Among these were mutations in and around HP1, suggesting that HP1 and HP2 do not share similar functional roles despite structural pseudosymmetry. Also mutations designed to increase interdomain hinge flexibility (P75G and E227G) did little to boost Glt_Ph_ activity, suggesting that hinges are sufficiently dynamic even in thermophilic glutamate transporters. Combining the three gain-of-function mutations, Y204L, A345V, and V366A produced a mutant with activity 12.7 ± 2.5 times higher than WT Glt_Ph_ (**Figures 1E, F** and **Supplementary Table 2**). Thus, subtle packing mutations in HP2 and the scaffold TM5 increase the transport rate by over an order of magnitude.

No other combinations produced further rate improvements, including those with the “humanizing” R276S/M395R mutations (**Supplementary Table 2**). To further boost transport, we considered the K290A mutation on HP1 (**Figure 1E**), which disrupts a salt bridge with E192 on the scaffold in the OFS and increases the transport domain dynamics ^37,42^. The K290A mutation by itself did not affect the transport rate. However, when combined with the Y204L/A345V/V366A mutant, it yielded our most active variant with 18 ± 1 times faster uptake rate than WT Glt_Ph_ and mean turnover time of ∼1 s (**Figures 1E, F** and **Supplementary Table 2**).

These robust, 10-20 fold, increases transport rate in a set of Glt_Ph_ variants that contain at most four mutations, are in line with similar efforts in enzymology ^68-74^.

### Rate-limiting steps of the transport cycle

To identify rate-limiting steps, we followed the timing of the transport domain movements between the OFS and IFS by smFRET, using total internal reflection fluorescence (TIRF) microscopy ^37,38,41,75^. Single cysteine mutations (N378C) were introduced within the transport domains of the Glt_Ph_ variants and labeled with self-healing fluorophores LD555P-MAL and LD655-MAL, and biotin-polyethylene glycol-maleimide. The labeled transporters were reconstituted into proteoliposomes and immobilized via a streptavidin-biotin bridge in microfluidic perfusion chambers enabling rapid buffer exchange (**Supplementary Figure 2**). We then monitored the relative movements of the donor- and acceptor-labeled transport domains within the trimeric transporters at 100 ms time resolution for a mean duration of ∼70 s before photobleaching occurred. We measured FRET efficiency (*E*_*FRET*_) of ∼0.4 when both transport domains were in the OFS. When one or both isomerized into the IFS, we observed increases of *E*_*FRET*_ to ∼0.6 and ∼0.9, respectively (**Supplementary Figure 2**).

In apo WT Glt_Ph_, when both the internal and external proteoliposome buffers contained no Na^+^ ions or l-Asp, the transport domains shuttled between the OFS and IFS with a mean frequency of 0.35 ± 0.01 s^-1^ (**Figure 2A**). When we replaced the external buffer with a buffer containing saturating concentrations of Na^+^ ions and l-Asp to establish the chemical gradients required for transport, we observed a dramatic reduction of the transport domain dynamics to a mean frequency of 0.04 ± 0.01 s^-1^ (**Figure 2B**). The dynamics of the Glt_Ph_ mutants showed similar overall features and trends. All variants exhibited comparatively fast dynamics under apo conditions with frequencies between ∼0.24 and 0.6 s^-1^ (**Figures 2A** and **Supplementary Figure 3**), which decreased to different extents under transport conditions, except in R276S/M395R Glt_Ph_ (**Figures 2B** and **Supplementary Figure 4**). In the presence of 10 mM blocker, D,L-threo-β-benzyloxyaspartic acid (D,L-TBOA), which is expected to suppress all transitions, we still observed a transition frequency of 0.03 ± 0.01 s^-1^ in both WT and mutant Glt_Ph_ proteins (**Supplementary Table 3)**. However, these transitions originated from ∼8-17 % of all molecules and constituted an apparent background. When this small sub-population was subtracted from the measured transition frequencies, the resulting transition frequency for the WT Glt_Ph_ at room temperature reduced to ∼0.01 s^-1^, on par with the transport turnover rate of 0.06 s^-1^ measured at 35 °C (**Supplementary Table 2**). Overall, these results are broadly consistent with the earlier measurements showing that the substrate-loaded transport domain is significantly less dynamic than the apo domain and that its movements occur with rates comparable to the uptake rates ^37,38^.

**Figure 2:**
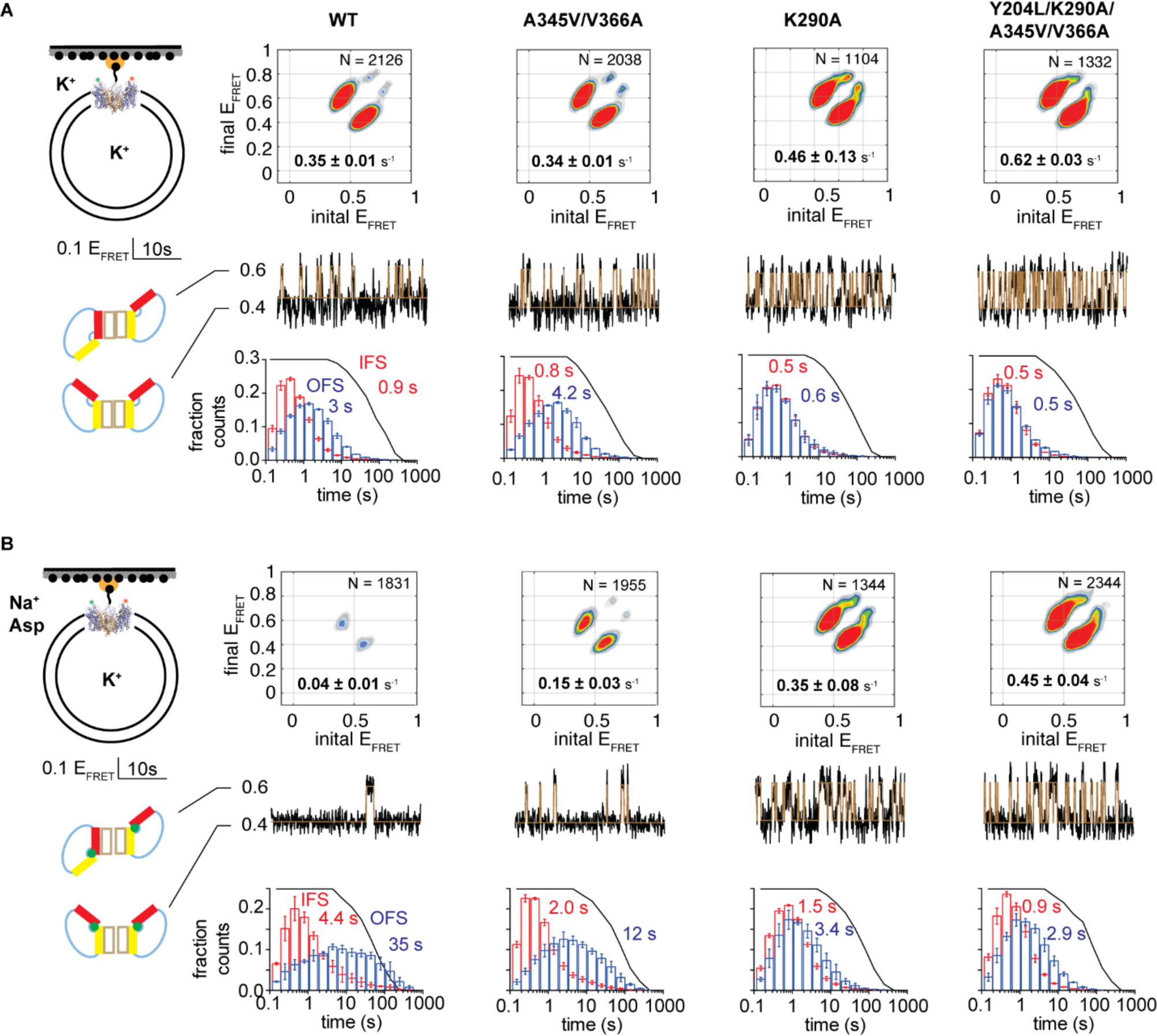
Transport domain dynamics under apo (A) and transport (B) conditions. Schematic representations of the experimental conditions are shown on the far left. Data are shown for WT Glt_Ph_ and select gain-of-function mutants, as indicated above the panels. Transition density plots (top of each panel) show the frequency of transitions between the *E*_*FRET*_ values. The number of trajectories analyzed (*N*) and the population-wide mean transition frequency is shown on the panels. Representative 50 s sections of single-molecule *E*_*FRET*_ trajectories (middle) with raw data in black and idealizations in brown. Scale bar and the conformational states corresponding to the low and intermediate *E*_*FRET*_ values are shown to the left of the panels. Dwell-time distributions for the OFS in blue and the IFS in red (bottom). Mean dwell times are shown on the panels in corresponding colors. Black lines represent photobleaching survival plots normalized from 1 to 0. Shown are means and standard errors over at least three independent repeats. *See also Supplementary Figures 2 –4.*

Correlated increases in dynamics and l-Asp uptake rates were observed in several mutants (**Figure 3A**). For instance, Y204L/A345V/V366A Glt_Ph_ showed that the mean transition frequency and the uptake rate increased by 12 and 13-fold, respectively. In contrast, the K290A and K290A/Y204L Glt_Ph_ mutants showed a ∼30-fold increase in dynamics but no significant changes in activity (**Figure 3A**). Consistent with our *K*_*m*_ measurements, we found that some mutations, including Y204L and K290A, had no significant effect on the affinity of the transporters for l-Asp. In contrast, the HP2 mutations A345V and V366A and the R276S/M395R mutant reduced affinity ∼10, 150, and 40 times, respectively (**Figures 3A** and **Supplementary Figure 5**). Most remarkably, combining mutations that dramatically increase the transition frequency (K290A and K290A/Y204L) with those reducing l-Asp affinity (A345V and V366A) yielded the fastest Glt_Ph_ variants (**Figure 3A**). Notably, l-Asp binding to the R276S/M395R mutant is several-fold faster than to the WT transporter ^57^. Therefore, it is likely that these and other mutations decrease l-Asp affinity by increasing the dissociation rate. Taken together, our data suggest that the translocation of the substrate-bound transport domain is the rate-limiting step of the cycle, but that substrate release can become rate-limiting in more dynamic mutants (**Figure 1B**).

**Figure 3:**
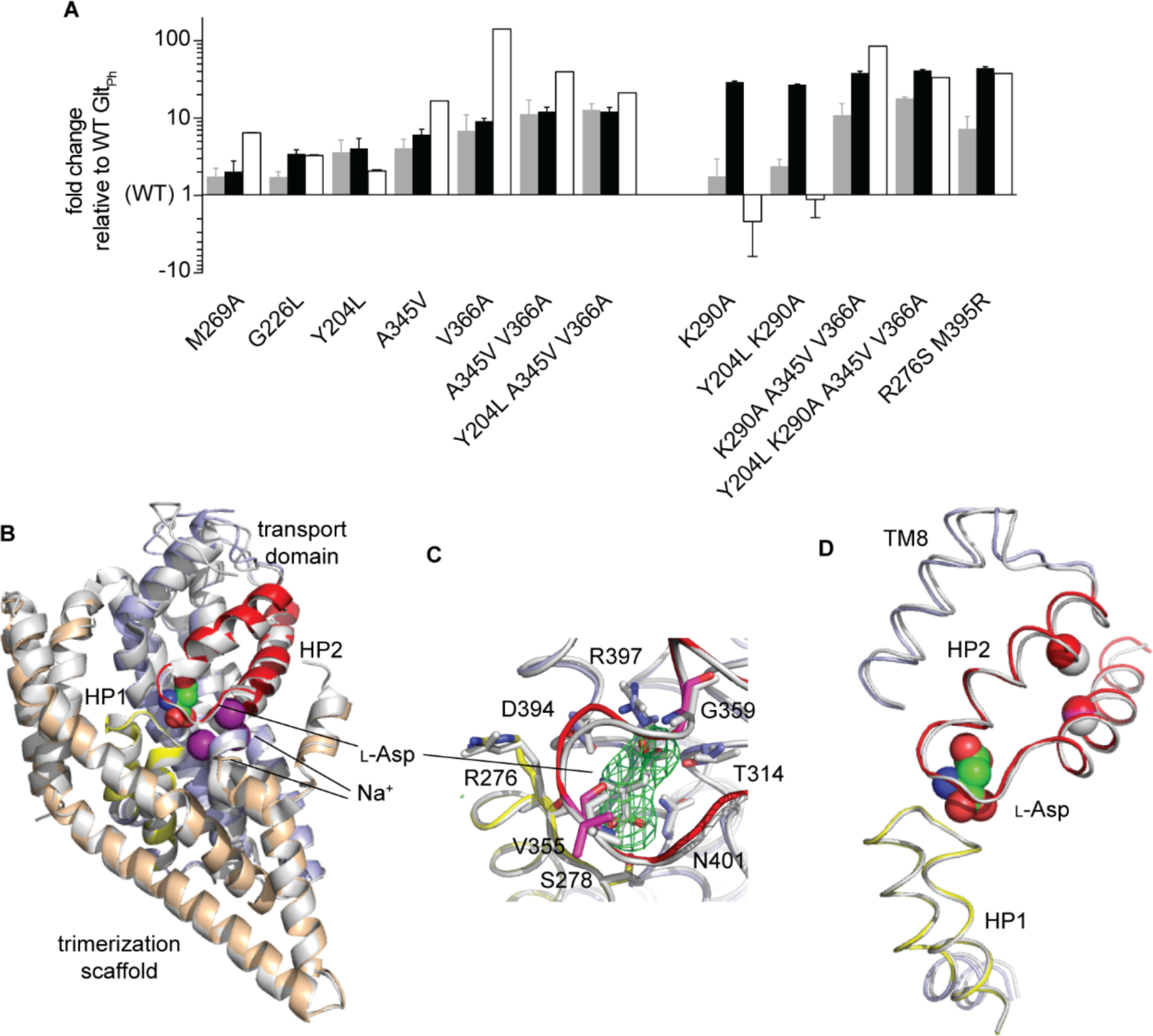
Correlations between activity, dynamics and substrate affinity. (**A**) Changes in the initial rates of substrate uptake (gray bars), transport domain dynamics (black bars), and l-Asp dissociation constant (white bars) of the mutant Glt_Ph_ variants relative to the WT transporter. The transition frequencies were measured under non-equilibrium transport conditions, and frequencies obtained in the presence of _D,L_-TBOA were subtracted from the data. Error bars represent standard errors for at least three independent measurements. (**B**) Cartoon representation of the crystal structure of Y204L/A345V/V366A Glt_Ph_ in the presence of saturating concentrations of Na^+^ ions and l-Asp (colored as in Figure 1A; PDB accession number 6V8G) superimposed onto the structure of WT Glt_Ph_ in the OFS (gray, PDB accession number 2NWX). l-Asp and Na^+^ are shown as spheres and colored by atom type. (**C**) Close-up of the substrate-binding site. Green mesh represents an omit electron density map for l-Asp contoured at 4*σ*. Substrate-coordinating residues are shown as sticks. (**D)** Superimposition of the mutant (colors) and the WT (gray) hairpins aligned on HP1. *See also Supplementary Figure 5 and Supplementary Tables 3 and 5.*

HP2 packing mutations A345V and V366A are striking because they both increase the transport domain dynamics and reduce l-Asp affinity. The altered binding was unexpected because the residues are located ∼14 Å away from the binding site (**Supplementary Figure 1**). The crystal structure of the Y204L/A345V/V366A mutant at ∼3.4 Å resolution showed the transporter in an OFS conformation nearly identical to that of the WT (**Figures 3B, D, Supplementary Table 4**). The electron density for l-Asp was visible, and the amino acid coordination was unaltered (**Figure 3C**). Interestingly, the unusually high B-factors of residues in HP2 and the adjacent part of TM8 suggested positional disorder in this region (**Supplementary Figure 5**). One notable difference from the WT structure was the position of the loop connecting TMs 3 and 4. The electron density was unresolved for the bulk of the loop, but a stretch of nine modeled N-terminal residues ran toward the top of the transport domain, instead of crossing over the HP2 surface as in WT Glt_Ph_ (**Supplementary Figure 5**). A similar loop conformation was seen in the crystal structure where the transport domain adopted an intermediate position between the OFS and the IFS ^67^. It remains unclear whether the observed loop conformations are consequences of different crystal packing or have functional implications ^76,77^. Overall, the mutations do not change the structure of HP2 or the binding site but might affect the local dynamics. If so, the reduced affinity could be due to the increased entropic penalty incurred upon substrate binding to a more dynamic apo protein or a less effective water exclusion from the binding site.

### The predominant mode of the transport domain dynamics

To study the energy barrier controlling the rate of substrate translocation, we collected smFRET recordings for Glt_Ph_ variants showing increased dynamics when both the internal and external proteoliposome buffers contained equal, saturating concentrations of Na^+^ ions and l-Asp. Under these equilibrium conditions, the substrate-bound transport domains shuttled between the OFS and IFS with the overall dynamics similar to those observed under the non-equilibrium transport conditions (**Supplementary Figure 4 and 6**). As observed in earlier studies ^38^, the WT Glt_Ph_ dynamics were highly heterogeneous, with the OFS and IFS dwell-times ranging from shorter than a second to hundreds of seconds (**Figure 4A**). At least three exponentials were required to fit the cumulative survival plots (**Supplementary Figure 6**), yielding lifetimes for the OFS of ∼3, 20, and 85 s, with each component contributing equally to the fit (**Supplementary Figure 6**). The IFS survival plots showed similar features, except that the short lifetime component was more prevalent (**Supplementary Figure 6**). The longest OFS dwells were shorter in more dynamic mutants, and this change was most dramatic in variants carrying the K290A and R276S/M395R mutations (**Figure 4A** and **Supplementary Figure 6**).

**Figure 4:**
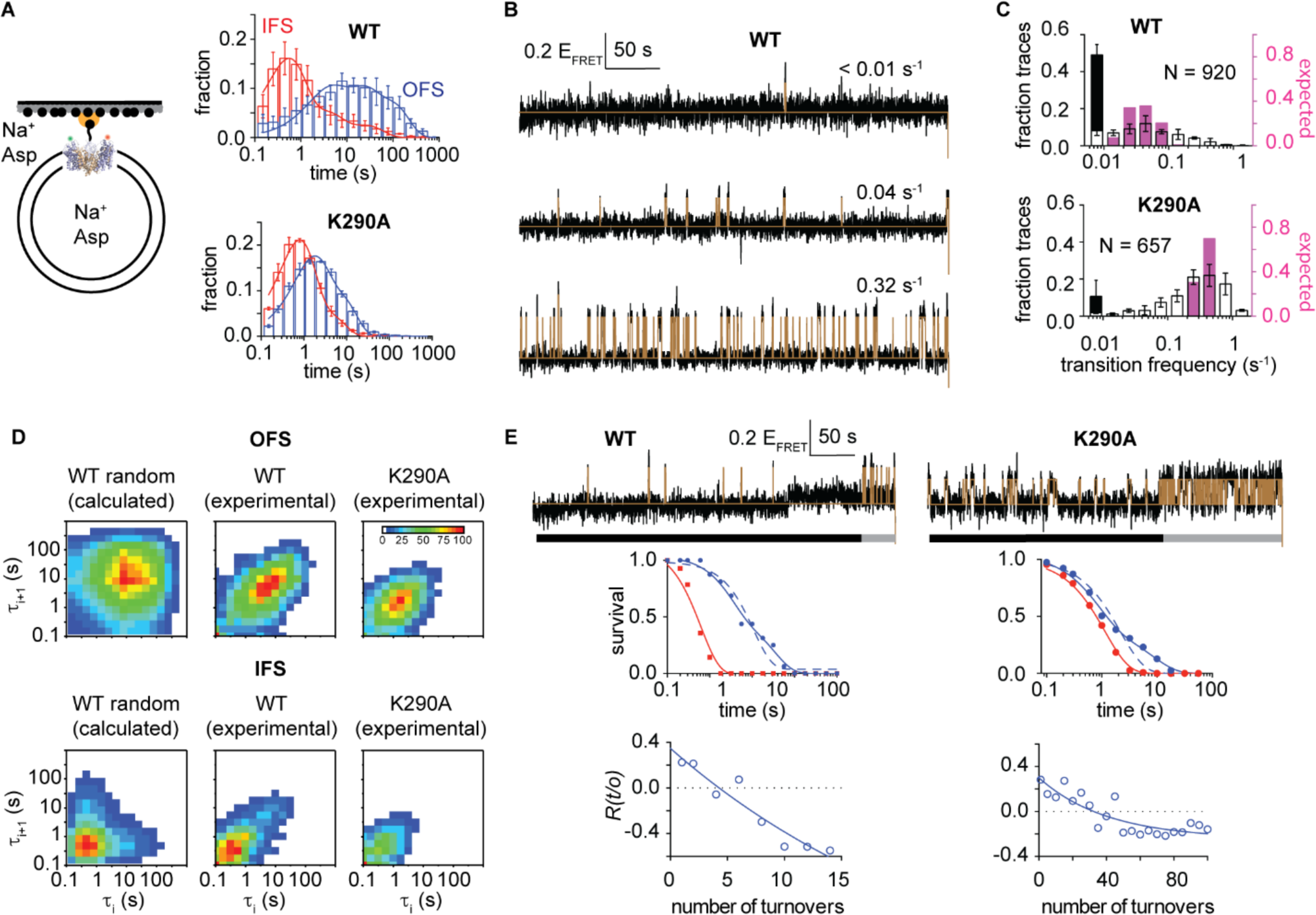
Kinetic heterogeneity of the transport domain dynamics. (**A**) Dwell-time distributions of the OFS (blue) and the IFS (red) for WT (top) and K290A (bottom) Glt_Ph_ observed under equilibrium conditions in saturating Na^+^/l-Asp (left). Lines are fits to three exponentials. (**B**) Representative *E*_*FRET*_ trajectories of WT Glt_Ph_ showing different transition frequencies. Raw data are in black, and idealizations are in brown. Scalebar is above the trajectories. (**C**) Distribution of the WT (top) and the K290A Glt_Ph_ (bottom) molecules with different mean transition frequencies (white bars). *N* is the number of trajectories longer than 90 s used in the analysis. The stacked black bars are fractions of trajectories without transitions. Pink bars show the expected binomial distribution if all trajectories shared the mean transition frequency of 0.04 s^-1^ for the WT and 0.34 s^-1^ for the K290A. (**D**) 2D-histograms of the consecutive dwell lengths in the OFS (top) and the IFS (bottom). From left to right: calculated distribution of dwell times randomly selected from the distributions in panel A; measured distribution for WT Glt_Ph_ and the K290A mutant. (**E**) Representative trajectories of the WT (left) and the K290A (right) Glt_Ph_ molecules showing switching between dynamic modes (top). Black and grey bars under the trajectories indicate apparent slower and faster dynamic modes. Survival plots for the OFS and the IFS (middle). Solid lines are fits to a single (IFS) and double (OFS) exponentials. Dashed lines (OFS only) are rejected fits to single exponentials. Autocorrelation plots of the sequential dwell durations (bottom). Solid lines are fits to single exponentials to guide the eye. *See also Supplementary Figures 6-9*.

Strikingly, some smFRET traces for the WT transporter showed no or rare transitions, while others featured sustained dynamics (**Figure 4B**). When we calculated transition frequencies for the individual traces lasting longer than 90 s before photobleaching, we obtained a distribution spanning over two orders of magnitude (**Figure 4C**). This distribution was significantly broader than would be expected if dwells occurred randomly, and all such molecules had the same intrinsic dynamics (**Figures 4C**). Much broader than expected transition frequency distributions were also observed for all mutants exhibiting increased dynamics, such as K290A Glt_Ph_ (**Figures 4C** and **Supplementary Figure 7**). These data indicate that the dynamics of Glt_Ph_ and Glt_Ph_ mutants are characterized by static disorder ^78^, i.e., distinct kinetic behaviors persist for periods of time comparable to or longer than the observation window. Consistent with this notion, we observed correlated lengths of the consecutive dwells in all Glt_Ph_ variants (**Figure 4D** and **Supplementary Figure 8**). Moreover, the majority of the individual traces showed intrinsically homogeneous dynamics. Their OFS and IFS survival plots were fitted well by single exponentials (**Supplementary Figure 9**), and the dwell lengths varied randomly around the means as revealed by the flat autocorrelation functions ^79^ (**Supplementary Figure 9**). Only in a minor fraction of molecules, between 5 and 20 % for both WT and mutant Glt_Ph_, were the survival plots better fit by two exponentials. Notably, in a subset of these molecules, we observed an apparent switching from one dynamic mode to another, which resulted in autocorrelation functions indicative of temporal segregation of the similar dwells (**Figures 4E** and **Supplementary Figure 9**). These results are generally in agreement with our earlier recordings that showed that the transporters sampled dynamic states and also long-lasting quiescent states (Akyuz, 2015), which we attributed to distinct off-pathway “locked” conformations. In our current recordings, which are approximately three times longer due to the improved stability of the self-healing fluorophores ^80,81^, we observe that the previously described “locked” states can indeed transition into the IFS, but with low frequency.

Each Glt_Ph_ transporter with intrinsically homogeneous dynamics can be described by a pair of characteristic OFS and IFS lifetimes. When plotted on a 2D histogram, they visualize the kinetic heterogeneity of the entire population (**Figure 5A**). For WT Glt_Ph_, the histogram showed that “typical” molecules had long OFS lifetimes, ranging from ∼10 to 100 sec, and short IFS lifetimes between ∼0.5 and 2 s (**Figure 5A**). The same characteristic kinetic mode prevailed in all of the mutants. Some mutants, most strikingly A345V and V366A Glt_Ph_, also had an increased fraction of transporters with long IFS and very short OFS lifetimes (**Figure 5B**). At present, the functional significance of these distinct kinetic behaviors is unclear. Consistent with this observation, recently established single-molecule transport assays have also revealed that WT Glt_Ph_ exhibits a broad kinetic heterogeneity, with individual transporters showing turnover times between seconds and hundreds of seconds (Ciftci et al., 2020, Science Advances, in review after revisions). Thus, the multiple kinetic modes of the conformational dynamics appear to parallel the multiple transport activity modes.

**Figure 5:**
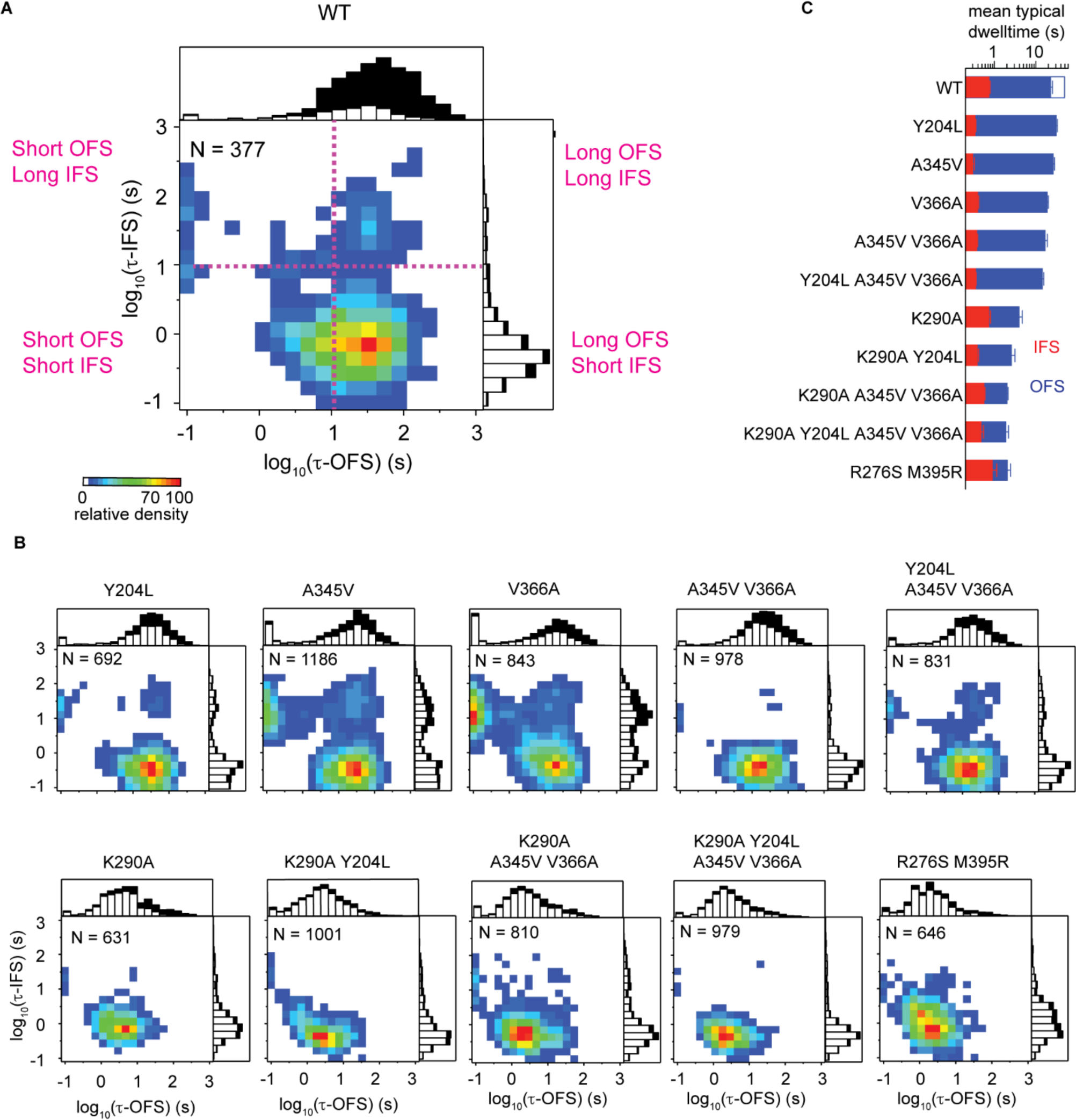
Distributions of the OFS and the IFS lifetime pairs for individual molecules. (**A** and **B**) 2D-histograms of lifetime pairs obtained for trajectories exhibiting single exponential behavior. *N* is the number of traces used. Scale bar shows relative density normalized by the number of molecules. Above and to the right of each panel are stacked histograms of, respectively, the OFS and the IFS lifetimes of the analyzed molecules (open bars) and the molecules showing no transitions before photobleaching (black bars). Data from three independent measurements were combined for presentation. (**C**) The mean lifetimes of the molecules falling within 30 % of the most populated bins of the 2D-histograms (yellow to red). The open blue bar represents the calculated mean OFS lifetime for WT Glt_Ph_ (see main text for details). Shown are means and standard errors of at least three independent measurements.

In this context, we note that the observed kinetic heterogeneity is unlikely to be due to post-translational modifications or differences in the lipid environments between individual vesicles, because it is reduced under apo conditions (**Supplementary Figure 10**). Instead, we hypothesize that the kinetic heterogeneity exhibited by Glt_Ph_ is indicative of a highly rugged energy landscape and the existence of long-lived conformations with distinct kinetic properties. Kinetic heterogeneities have been reported for many systems ^79,82-87^. However, they might be particularly pronounced in Glt_Ph_ because it is a thermophilic protein that might encounter high enthalpic barriers at ambient temperatures. To gain insight into the nature of the most commonly used activation barrier during transport domain motions, we set out to perform TS analyses on the predominant dynamic mode shared by the WT protein and the dynamic mutants.

### Transition state structure

While the macroscopic rates of the transport domain translocation are slow, translocation of the transport domain is faster than the resolution of our smFRET recordings (10-100 ms). Correspondingly, intermediate positions of the domain, manifesting in intermediate *E*_*FRET*_ values, are not readily observed. Therefore, diffusive movements of the transport domain across the membrane are fast, and the long dwell times in the OFS and IFS arise from relatively high energy barriers along the translocation trajectory. Because the transport domain undergoes a concerted, rigid body movement, we applied LFER analysis to infer the structure of the TS in terms of the transport domain position along the trajectory from the OFS to the IFS. LFER analysis correlates changes of the TS free energy relative to the equilibrium end states (i.e., the height of the free energy barrier) to the changes of the free energy difference between the end states in response to perturbations, such as mutations or ligand additions or removals (**Supplementary Figure 11**). The relative free energies of the TS and the end states are inferred from the forward and reverse reaction rate constants. If the TS resembles the OFS, mutations or ligands will change its free energy as much as the free energy of the OFS. If so, the change in free energy of the TS with respect to the OFS, ΔΔG^#^_OFS to TS_, will be near zero. Thus, the height of the energy barrier crossed during the transition from the OFS to the IFS, and the corresponding forward rate constant, *k*_*OFS to IFS*_, will be unaltered. In contrast, the height of the free energy barrier crossed during the reverse transition from the IFS to the OFS, ΔΔG^#^_IFS to TS_ will change as much as the free energy of the IFS relative to the OFS, and the reverse rate constant, *k*_*IFS to OFS*_, will change accordingly (**Supplementary Figure 11**). If, however, the TS resembles the IFS, the above relationships will be reversed. We will observe the forward rate constants, *k*_*OFS to IFS*_, that change in response to the perturbations, and the reverse rate constants, *k*_*IFS to OFS*_, that do not (**Supplementary Figure 11**). Already a qualitative comparison of the smFRET recordings of WT Glt_Ph_ in the absence and presence of the substrate and Na^+^ ions shows that the ligands mostly affect the duration of the OFS dwells (*k*_*OFS to IFS*_). By contrast, the IFS dwells (*k*_*IFS to OFS*_) are unchanged, hinting that the transition state might resemble the IFS (**Figure 11C**).

Quantitatively, the linear relationship between the equilibrium and the TS free energies is expressed as follows ^30-32^:

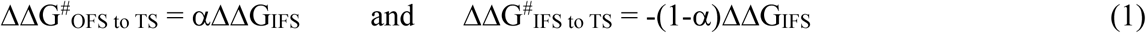

where ΔΔG_IFS_ is the change of the free energy of the IFS relative to the OFS. The Leffler α approaches 0 or 1 when the TS resembles the OFS or the IFS, respectively (**Figure 6A**, *left and right*). Notably, if the transport domain in the TS assumes an intermediate position between the OFS and IFS, we will obtain different α values depending on the location of the perturbation (**Figure 6A**, *middle*). Mutations of residues involved in the same interactions in the TS as in the OFS or IFS will yield α values of 0 and 1, respectively. Mutating residues involved in interactions distinct from both will yield intermediate values of α.

**Figure 6:**
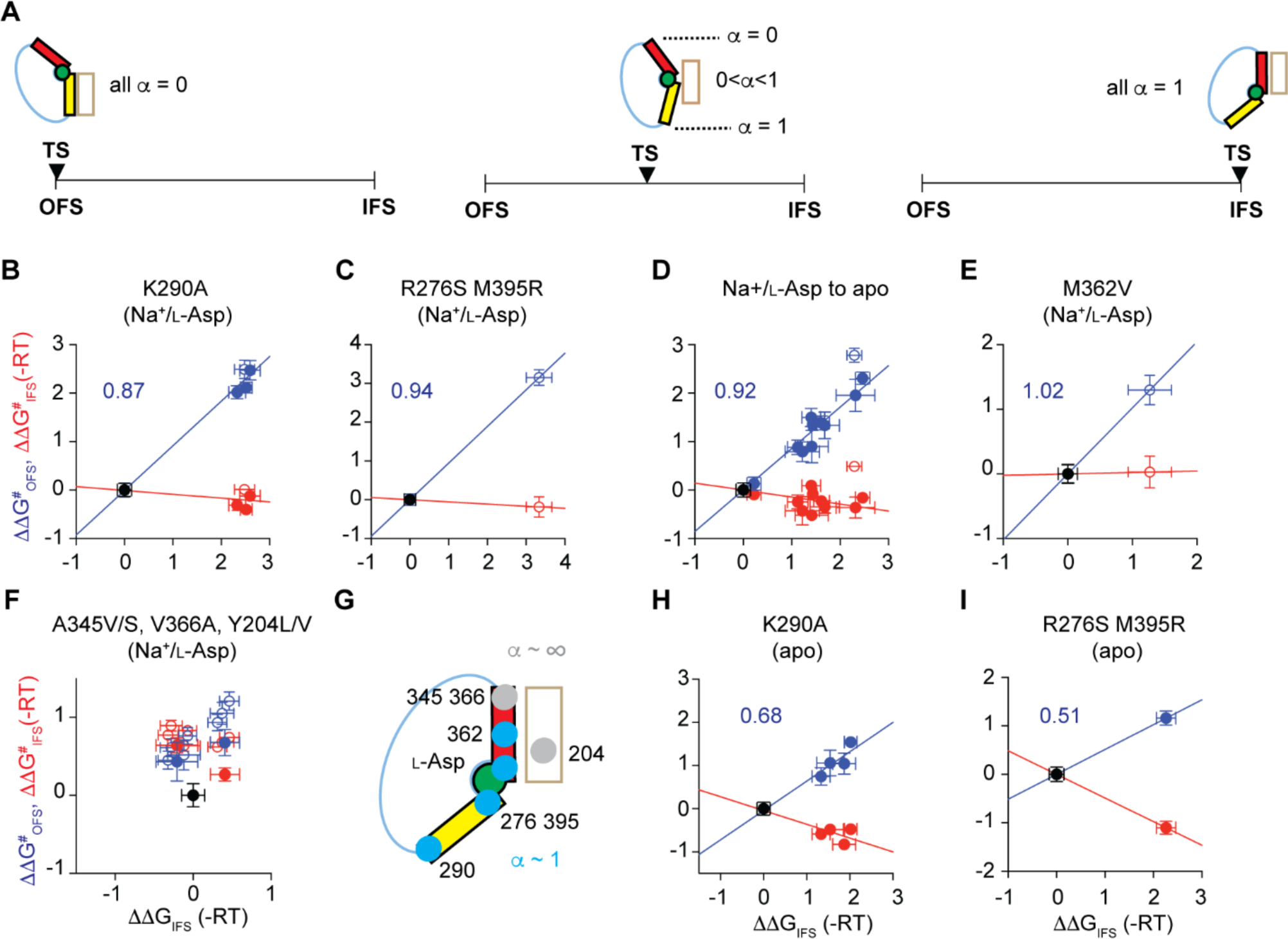
Transition state analysis. (**A**) Schematic representation of the expected Leffler α values if the transition state is structurally similar to the OFS (left) or the IFS (right) or if it assumes an intermediate structure (middle). (**B-F, H, I**) LFERs with free energy changes plotted in units of -RT. The activation free energies of the OFS to IFS transitions (blue) and the IFS to OFS transitions (red) were calculated by subtracting the energies of the reference states (black, at the origin) from the energies of the mutated protein variants. The K290A mutation was introduced into the WT, Y204L, A345V/V366A, and Y204/A345V/V366A backgrounds, either substrate-bound or apo (**B** and **H**, respectively). R276S/M395R and M362V mutations were introduced into the WT background (**C, E**, and **I**). For the transition from Na^+^/l-Asp-bound to apo state, the free energies measured for the transporters in the presence of Na^+^ and l-Asp were subtracted from those measured for the apo transporters (**D**). Mutations at A345, V366 and Y204 sites and their combinations were introduced within the WT and K290A backgrounds (**F**). LFERs for the perturbations introduced within the WT background use back-calculated WT transition rates (see Main text, open symbols). (**G**) Schematic summary of sites where perturbations led to changes of the transition state energy that scaled with the IFS energy (blue) and where they affected only the transition state (grey). *See also Supplementary Figure 10 and 11.*

Our complement of mutants allowed us to test the effects of perturbations along the scaffold-facing transport domain surface from the cytoplasmic base of HP1 to the extracellular base of HP2. For each, we estimated the rate constants, *k*_*OFS to IFS*_ and *k*_*IFS to OFS*_, from the inverse of the mean OFS and IFS lifetimes of the molecules in the predominant dynamic mode (**Figure 5B** and **Supplementary Figure 10**). First, we examined the effects of the K290A mutation at the base of HP1 (**Figure 1A**) introduced into the Y204L, A345V/V366A, and Y204L/A345V/V366A background mutants, because their characteristic OFS lifetimes were well determined (**Figure 5B**). We observed that the K290A mutation shortened the lifetimes of the OFS in these backgrounds (increased *k*_*OFS to IFS*_) by 9 ± 2 times on average but had little effect on the IFS lifetimes (**Figure 5C**). The free energies of the IFS and the TS decreased similarly by ∼2.5 RT relative to the OFS with the Leffler α of 0.87 ± 0.06 (**Figure 6B**). Based on these results, we conclude that the salt bridge between K290 and E192 in the scaffold domain is already broken in the TS of Glt_Ph_ molecules. The effect of the K290A mutation on WT Glt_Ph_ was similar to the other variants with only the OFS lifetime affected (**Figure 5C**). However, we were unable to measure the WT Glt_Ph_ OFS lifetime directly, because it was comparable to the fluorophore lifetime (**Figure 5A**). We therefore assumed that the mutation should shorten the OFS lifetime to the same extent in the WT as in the other backgrounds and back-calculated the WT OFS lifetime to be ∼49 s from the lifetime of K290A Glt_Ph_. We used this corrected value in comparisons with other mutations.

The R276S/M395R mutations at the tip of HP1 (**Figure 1A**) also shifted the equilibrium toward the IFS, and the free energy of the TS and the IFS decreased similarly, with Leffler α of 0.94 (**Figure 6C**). We obtained similar results when we compared the translocation rates of all mutants in the presence of the saturating concentrations of Na^+^ ions and l-Asp, and under apo conditions, yielding mean Leffler α of 0.92 ± 0.05 (**Figure 6D**). An HP2 mutation M362V also showed a small but similar decrease of the relative free energies of the TS and the IFS (**Figure 6E**). R276 in HP1 and M395 in TM8 face the extracellular and the cytoplasmic solutions in the OFS and the IFS, respectively, and are poised to interact with different parts of the scaffold domain. Furthermore, HP2 undergoes a conformational change in the apo Glt_Ph_, which alters the interface between the transport and scaffold domains in the IFS, but not in the OFS ^54^. Thus, our analysis suggests that HP1, HP2, and TM8 in the TS, form interactions that are similar to those in the IFS. These findings are consistent with the transport domain completing most of the OFS to the IFS movement prior to overcoming the principal activation barrier needed to achieve the IFS.

Finally, we examined mutations of A345 and V366 residues near the extracellular base of HP2 and Y204 in TM5 kink of the scaffold domain (**Figure 1A**). Surprisingly, multiple mutations at these sites led to the increases of both *k*_*OFS to IFS*_ and *k*_*IFS to OFS*_ when introduced into either the WT or the K290A Glt_Ph_ backgrounds (**Figure 6F**). Increased rates of both forward and reverse reactions are a hallmark of mutations that specifically stabilize the TS ^31,88^. Thus, the TS structurally resembles the IFS but is sufficiently distinct so that mutations at Y204, A345, and V366 sites affect the two states differentially (**Figure 6G**). Notably, when we constructed LFERs for the apo transporters, we observed Leffler α values of ∼0.7 and ∼0.5 for K290A and R276S/M395R mutants, respectively (**Figures 6H – I**). These results suggest that disrupting the domain interface in the OFS might become a kinetically more critical step when the barrier near the IFS diminishes ^31,89,90^.

## Discussion

To probe the structure of the rate-limiting TS for transport domain movement in Glt_Ph_, we first identified gain-of-function mutations by comparing sequences of homologues from hyperthermophilic, thermophilic, mesophilic, and psychrophilic bacteria. Our functional assays and smFRET analyses of these Glt_Ph_ mutants showed that the increased frequency of transitions between the OFS and the IFS of the l-Asp-bound transport domain, and the decreased l-Asp affinity, lead to faster uptake rates. Synergistic changes of the dynamics and affinity were observed in combination mutants and also in single HP2 mutants. Interestingly, the populations of the OFS and the IFS in the HP2 mutants remained similar in the absence and presence of l-Asp and Na^+^ ions (**Supplementary Table 5**). This finding suggests that their OFS and IFS have similar affinities for the substrate as is also the case for the WT transporter ^91^. Our analyses further show that mutations in HP2 affect substrate binding similarly in the OFS and the IFS and suggest that HP2 plays a similar gating role in the two states. Crystal structures of the WT Glt_Ph_ in the IFS pictured HP2 immobilized at the domain interface unable to act as a gate ^49,54,67^. In contrast, structures of the R276S/M395R Glt_Ph_ mutant, the archaeal homologue Glt_Tk_ ^92^ and the human homologue ASCT2 ^38,56^ showed so-called “unlocked” conformations, in which the transport domain leans away from the scaffold, providing space for HP2 to open. Our current results support the “one gate” model ^56^, whereby HP2 gates substrate binding in both the OFS and the unlocked IFS. Collectively, these data suggest that the conformationally flexible HP2 ^34,50-55^ serves as a master-regulator of substrate binding, translocation, and release.

SmFRET recordings under equilibrium conditions in the presence of saturating Na^+^ ions and L-Asp showed that the majority of Glt_Ph_ molecules resided in the OFS, consistent with earlier studies ^37,38,41,42,45,47^. Long OFS dwells were interspersed by brief excursions into the IFS, but long IFS dwells were also observed in some molecules and sustained dynamics were seen in others. Overall, we found that the mean transition frequencies vary widely between the individual molecules in both the WT transporters and the more dynamic mutants. Such behavior is best described in terms of multiple dynamic modes, whereby molecules show long-lasting differences in the kinetics of the observed transitions. At present, the structural origins of the heterogeneity are not clear. It might arise from distinct protein conformations, but it is also possible that distinct lipid interactions or other effects account for the sustained differences in dynamics.

LFER analysis on nearly all introduced perturbations suggested that the TS for transport domain movement structurally resembles the IFS. Hence, once the transport domain is dislodged from the scaffold in the OFS, it might take multiple attempts to achieve the IFS conformation before successfully crossing the high-energy barrier. It is also interesting that subtle packing mutations in HP2 and the scaffold affect both forward and reverse translocation rates. Moreover, an altered structure of the domain interface was observed in the crystal structure of the dynamic R276S/M395R Glt_Ph_ mutant in the IFS^38^. Thus, we hypothesize that exploring the local conformational space in the IFS-like TS to achieve a stable configuration of the domain interface might be rate-limiting.

The structural similarity between the TS and the IFS suggests that molecules that bind at the interface of the transport domain and the scaffold with higher affinity for the IFS than for the OFS would also bind tighter to the TS. Such molecules would reduce the height of the rate-limiting energy barrier of the cycle and work as activators or positive allosteric modulators (PAMs) (**Figure 7A**). In contrast, molecules that bind tighter to the OFS would serve as allosteric inhibitors. The degree to which PAMs can accelerate the transport cycle would depend on how much tighter they bind to the TS and the IFS compared to the OFS and on the height of other, unaffected barriers (**Figures 7B, C**). By a similar logic, if the return of the apo transporter into the OFS were rate-limiting with the main energy barrier near the OFS, small molecules that bind tighter to the OFS than the IFS would serve as PAMs.

**Figure 7:**
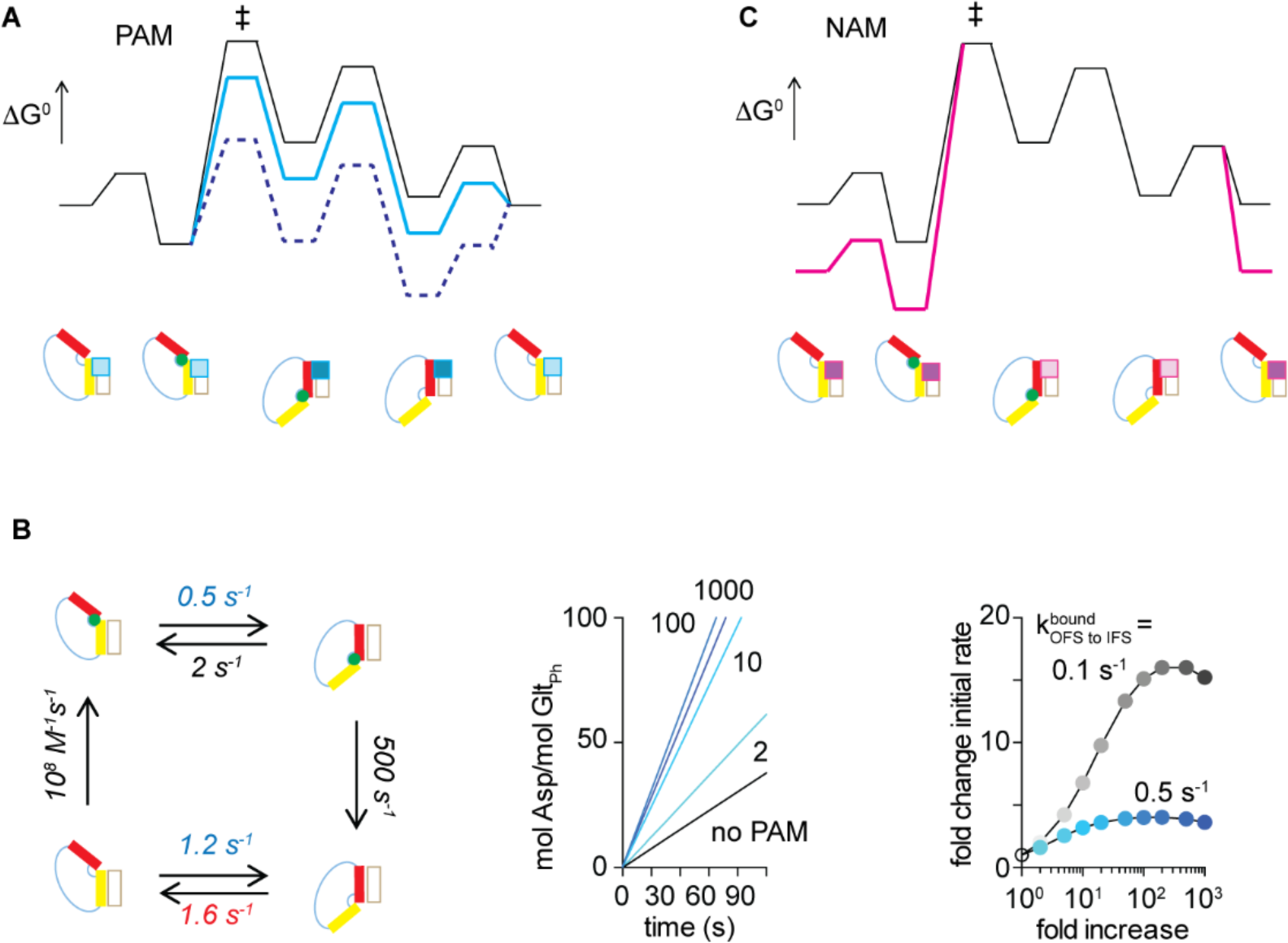
Proposed model for allosteric modulation. (**A, C**) Simplified free energy diagram of the transport cycle (black) and its modulation (colors) by PAMs (**A**) or NAMs (**C**). Cartoon representations of the low-energy states are below the diagrams. Substrate binding to the apo OFS precedes isomerization into the IFS via the highest-energy TS (‡) that structurally resembles the IFS. Substrate release and recycling into the OFS complete the cycle. PAMs, shown as squares in the cartoons of the states, bind with higher affinity to the IFS (dark cyan squares) than to the OFS (light cyan squares). Therefore, they stabilize all IFS-like states, including the TS, and smoothen the energy landscape (cyan line). PAMs that bind too tightly to the IFS may become inhibitory, as apo OFS becomes a high-energy state (dotted blue line). NAMs bind tighter to the OFS (**C**, dark magenta squares) than to the IFS (light magenta squares) and increase the ruggedness of the landscape (magenta line). (**B**) Simulations of the transport rates in the absence and presence of PAMs. The left panel shows the transport cycle with the used rate constants (**Methods**). Middle and right panels show simulated uptake and fold increase of the initial rates, respectively, in the presence of PAMs that increase *k*_*OFS to IFS*_ by the indicated number of folds (from cyan to dark blue). Simulations on cycles with 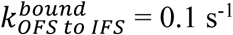 illustrate higher PAM potency (grey).

PAMs of human EAATs offer therapeutic opportunities to treat a plethora of conditions associated with glutamate-mediated excitotoxicity in the central nervous system ^93^. Recently developed compounds boost EAAT2 transport by ∼2-fold and provide neuroprotection (Falcucci et al., 2019; Kortagere et al., 2018). Their mechanism is not yet understood but some of their features agree with the concepts developed here. Specifically, the identified PAMs are thought to bind preferentially to the interface of the transport and scaffold domains in the OFS ^94,95^, consistent with the return of the apo transport domain into the OFS being the rate-limiting step of the cycle ^96^. Moreover, chemically similar compounds act as activators or as inhibitors ^95^ as predicted by our model.

Approximately 50 % of solute carriers are implicated in disease in humans, but only a few are drug targets ^97^. Our approach conceptualizes how the rate-limiting transition state of a transporter can be characterized structurally, informing on potential druggable sites to develop allosteric modulators.

## Online Methods

### Bioinformatics analysis

The workflow of the bioinformatics approach is in **Supplementary Figure 1**. Organism taxa, optimal growth temperatures, and temperature limits for growth were extracted from Bac*Dive* ^98^. Sequences of 20,000 prokaryotic homologues of Glt_Ph_ were collected using BLAST ^99^ and aligned using MAFFT 7.0, applying the highest gap penalty ^100^. Multiple sequence alignments (MSA) were manually adjusted to optimize the alignment of the secondary structure elements and to remove sequences that were lacking one or more transmembrane helices. Sequences with over 90 % amino acid sequence identity were clustered using USEARCH ^101^, and one random sequence per cluster was retained for subsequent analysis. Optimal growth temperatures and temperature limits were assigned to the sequences by matching GenInfo identifiers with genus and species annotations in Bac*Dive* database. Sequences without a match were excluded from further analysis. The remaining sequences were sorted into four temperature groups: psychrophiles (growth temperatures below 21 °C), mesophiles (21 °C to 45 °C), thermophiles (46 °C to 65 °C) and hyperthermophiles (over 65 °C). All temperature groups, except mesophiles, were manually sorted to exclude sequences originating from organisms with growth temperature ranges spanning more than two temperature groups. This process yielded 128, 5811, 93, and 24 sequences from psychrophiles, mesophiles, thermophiles, and hyperthermophiles, respectively. The amino acid frequencies were calculated at each position in the MSA for all temperature groups using Prophecy 102 and using the Glt_Ph_ sequence as a reference. Combined frequencies were calculated for amino acids classified by their physicochemical properties (A+V, G, S+C+T+N+Q, P, D+E, H+M, Y+W, F+L+I, K+R) or side chain volumes (A+G+S, P+C+T+N+D, H+E+Q+V, Y+W+F, M+L+I+K+R). We used the outcomes of two-sample Kolmogorov-Smirnov tests to determine whether amino acid distributions at each position of the MSA were statistically different between pairs of temperature groups. Because no position showed significant distribution changes across all temperature groups, the results of every pairwise comparison in the Kolmogorov-Smirnov test were weighted to reflect the temperature gap between the groups (i.e., multiplied by 3 for the comparison between the hyperthermophiles and psychrophiles, 2 for thermophiles and psychrophiles, *etc.*). A positive value of the sum of the weighted values (global difference score) for a given position in the MSA indicates a change of the amino acid distribution correlated to the optimal growth temperature. Among the amino acids with positive global difference score, mutation sites were selected according to the following criteria: (1) their side chains were not exposed to either aqueous solution or lipid bilayer; (2) they were in well-conserved regions; and (3) they were not directly coordinating l-Asp or Na^+^ ions. Additional residues in these areas were chosen to further perturb packing interactions (I85, L303, T322, I341, and M362). Differences in the crystal structures of WT Glt_Ph_ in the OFS and the IFS were used to guide mutagenesis of several residues in the inter-domain hinges (A70, P75, Q220, H223, G226, and E227) and TM2/5 kink (D48 and Y204). Glt_Ph_ residues were mostly mutated to those prevalent in mesophiles or psychrophiles, in most cases producing conservative mutations that lead to subtle volume changes or the removal of hydrogen bonds.

### Mutagenesis, protein expression, purification and labeling

Amino acid substitutions were introduced by site-directed mutagenesis (*Stratagene*) in a Glt_Ph_ variant with seven non-conserved surface-exposed residues replaced with histidines to increase expression yields (referred to as wild type) ^46^. For smFRET microscopy, substitutions were introduced within the Glt_Ph_ variant with additional C321A and N378C mutations, as previously described ^37^. Constructs were cloned in-frame into pBAD24 vector with a C-terminal thrombin cleavage site followed by octa-histidine tag as described previously ^46^ and verified by DNA sequencing. Plasmids were transformed into *E. coli* strain DH10B (*Invitrogen*), and protein expression was induced for 3 h at 37 °C by 0.1 % arabinose. Glt_Ph_ was purified from cell membranes, as described previously ^46^. Briefly, membranes were solubilized in buffer containing 20 mM Hepes/Tris, pH 7.4, 200 mM NaCl, 1 mM l-Asp and 40 mM *n*-dodecyl-β-D-maltopyranoside (DDM, *Anatrace*). Insoluble material was removed by ultracentrifugation at 100000 *g* for 1 h at 4 °C. Solubilized transporters were bound to nickel agarose (*Qiagen*) for 2 h at 4 °C. The resin was washed in the same buffer containing 1 mM DDM and 40 mM imidazole and proteins eluted with 250 mM imidazole. The tag was removed by overnight cleavage with thrombin at room temperature using 10 units per mg Glt_Ph_. The proteins were further purified by size exclusion chromatography on a Superdex 200 Increase 10/300 GL column in buffers of various compositions: 10 mM Hepes/Tris, pH 7.4, 200 mM NaCl, 1 mM l-Asp and 7 mM n-decyl-β-d-maltopyranoside (DM, *Anatrace*) for transport assays; 10 mM Hepes/Tris, pH 7.4, 50 mM NaCl, 0.1 mM l-Asp and 5 mM DM for crystallization; 10 mM Hepes/Tris, pH 7.4, 1 mM NaCl, 200 mM choline chloride, 1 mM DDM for binding assays; and 10 mM Hepes/Tris, pH 7.4, 200 mM NaCl, 1 mM l-Asp, 1 mM DDM for smFRET microscopy. When Glt_Ph_ proteins were purified for smFRET microscopy, all buffers were supplemented with 0.1 mM Tris(2-carboxyethyl)phosphine (TCEP). Glt_Ph_ purity was confirmed by SDS PAGE, followed by Coomassie Brilliant Blue R-250 staining. For smFRET studies, purified Glt_Ph_ variants at 40 µM were labeled using maleimide-activated LD555P-MAL and LD655-MAL dyes ^80,81^ and biotin-PEG_11_ (*Thermo Fisher Scientific*)) at final concentrations of 50, 100 and 25 μM, respectively, as previously described ^37^. Labeling efficiency was determined by spectrophotometry using extinction coefficients of 57400, 150000, and 250000 M^-1^cm^-1^ for Glt_Ph_, LD555P-MAL, and LD655-MAL, respectively. Excess dyes were removed on a PDMiniTrap Sephadex G-25 desalting column (*GE Healthcare*).

### l-Asp binding assays

Fluorescence binding assays were performed as described previously ^54^. Briefly, Glt_Ph_ was diluted to a final concentration of 2 μM in buffer containing 20 mM Hepes/Tris, pH 7.4, 200 mM choline chloride, 0.4 mM DDM, and 0.4 nM RH421 dye (*Invitrogen*) supplemented with 1 or 10 mM NaCl, as indicated. RH421 was excited at 532 nm, and emission was measured at 628 nm at 25 °C using a QuantaMaster fluorimeter equipped with a magnetic stirrer (*Photon International Technologies*). Fluorescence changes induced by additions of l-Asp aliquots were monitored until stable for ∼100 s, corrected for dilution, and normalized to the maximal fluorescence change. Binding isotherms were plotted and fitted to Hill equation using Prism (*GraphPad*). All measurements were performed in triplicate.

### Glt_Ph_ reconstitution into liposomes and transport assays

Glt_Ph_ variants were reconstituted into liposomes as described previously ^63^. Liposomes were prepared using *E. coli* polar lipid extract, egg yolk L-α-phosphatidylcholine and 1,2-dioleoyl-sn-glycero-3-phosphoethanolamine-N-(lissamine rhodamine B sulfonyl) (*Avanti Polar Lipids*) in a 3000:1000:1 (w/w/w) ratio. The dried lipid films were hydrated in buffer containing 20 mM Hepes/Tris, pH 7.4, 200 mM KCl, 100 mM choline chloride at a final concentration of 70 mM lipid by repeated freeze-thaw cycles. Liposomes were extruded through polycarbonate filters with a pore size of 400 nm (*Avanti Polar Lipids*) and destabilized with Triton X-100 (*Sigma*) at a detergent-to-lipid ration of 0.5:1 (w/w). Glt_Ph_ variants were added at final protein-to-lipid ratios of 1:2000 (w/w) and incubated for 30 min at 22 °C. Detergents were removed by repeated incubations with BioBeads SM-2 (*BioRad*). Liposomes were concentrated by ultracentrifugation at 100000 *g* for 1 h at 4 °C, subjected to three freeze-thaw cycles, and extruded through 400 nm polycarbonate filters (*Avanti Polar Lipids*). To initiate transport, liposomes were diluted 100-fold into a buffer containing 20 mM Hepes/Tris, pH 7.4, 200 mM KCl, 100 mM NaCl, 0.5 μM valinomycin and variable concentrations of ^3^H-l-Asp (Aspartic acid, L-[2,3-^3^H], *Perkin Elmer*). When necessary, the reaction mixtures were supplemented with cold l-Asp. At appropriate time points, 200 μl aliquots were removed and diluted into ice-cold quench buffer containing 20 mM Hepes/Tris, pH 7.4, 200 mM LiCl, 100 mM choline chloride, followed by rapid filtration using 0.22 μm nitrocellulose filters (*Wattman*). Filters were washed three times using a total of 8 ml quench buffer. The retained radioactivity was measured by scintillation counting in a LS-6500 counter (*Beckman-Coulter*). To determine l-Asp *K*_*M*_, concentration dependences were measured using 1 min incubation to achieve robust signals for all variants. Uptake time courses were measured at concentrations of l-Asp of 5 times over *K*_*M*_ for 90 s. All uptake experiments were performed at 34 °C. The background was determined in the reaction buffer lacking NaCl and subtracted from the measurements. The proteoliposome concentration was determined by measuring rhodamine fluorescence using excitation and emission wavelengths of 530 and 590 nm, respectively. The protein concentration was estimated by assuming similar reconstitution efficiencies for all Glt_Ph_ variants. The amount of l-Asp uptake was normalized per Glt_Ph_ monomer. Dose-response curves and initial time courses were plotted and fitted in Prism (*GraphPad*).

### Crystallography

Purified Y204L/A345V/V366A Glt_Ph_ mutant was concentrated to 3 mg/ml. Protein was mixed 1:1 with a reservoir solution containing 100 mM potassium citrate pH 4.4 – 5 and 12 – 17 % PEG400. The protein was crystallized at 4 °C by hanging drop vapor diffusion method. Crystals were cryo-protected in reservoir solution supplemented with 30 – 35 % PEG400. Diffraction data were collected at Advanced Light Source beamline 8.2.2. Diffraction data were indexed, integrated, and scaled using HKL2000 ^103^. Initial phases were determined by molecular replacement in Phaser ^104^ using 2NWX as the search model. The model was optimized by iterative rounds of refinement and rebuilding in Phenix ^105^ and Coot ^106^. Strict non-crystallographic three-fold symmetry was applied during refinement.

### smFRET microscopy and data analysis

Glt_Ph_ was reconstituted into liposomes as above with modifications. The 1,2-dioleoyl-sn-glycero-3-phosphoethanolamine-N-(lissamine rhodamine B sulfonyl) was omitted, and the lipid film was hydrated in buffer containing 20 mM Hepes/Tris, pH 7.4 and 200 mM KCl. Glt_Ph_ was added at a final protein-to-lipid ratio of 1:1000 (w/w) to maximize the number of liposomes containing one trimer. Liposomes were extruded through 100 nm polycarbonate filters (*Avanti Polar Lipids*). To replace internal liposome buffer, vesicles were subjected to three rounds of the following procedure. Proteoliposomes were pelleted by centrifugation for 40 min at 100000 *g* at 4 °C, resuspended in buffer containing 20 mM Hepes/Tris, pH 7.4, 200 mM NaCl or 20 mM Hepes/Tris, pH 7.4, 200 mM NaCl and 0.1 mM l-Asp, as required, and subjected to 3 freeze/thaw cycles.

All smFRET experiments were performed on a home-built prism-based total internal reflection fluorescence (TIRF) microscope constructed around a Nikon Eclipse *Ti* inverted microscope using passivated microfluidic imaging chambers ^75,107^. The samples were illuminated with a 532 nm laser (*LaserQuantum*). LD555P and LD655 fluorescence signals were separated using a T635lpxr dichroic filter (*Chroma*) mounted in a MultiCam apparatus (*Cairn*). Imaging data were acquired using home-written acquisition software and scientific complementary metal-oxide semiconductor (sCMOS) cameras (*Hamamatsu*).

LD555P-MAL/LD655-MAL-labeled Glt_Ph_ variants in proteoliposomes were surface-immobilized via a biotin-streptavidin bridge on PEG-passivated microfluidic imaging chambers functionalized with streptavidin. Imaging experiments were performed in 20 mM Hepes/Tris buffers at pH 7.4 containing 5 mM β-mercaptoethanol and an oxygen scavenger system comprised of 1 U/ml glucose oxidase (*Sigma*), 8 U/ml catalase (*Sigma*) and 0.1 % glucose (*Sigma*). Experiments under symmetric apo conditions were performed in buffer supplemented with 200 mM KCl using proteoliposomes loaded with buffer containing 200 mM KCl. Under non-equilibrium transport conditions, imaging buffer was supplemented with 200 mM NaCl and 0.1 mM l-Asp. For symmetric saturating Na^+^/l-Asp conditions, the internal liposome and imaging buffers contained 200 mM NaCl and 0.1 mM l-Asp. To inhibit protein dynamics, liposomes containing 200 mM NaCl were incubated in imaging buffer contained 200 mM NaCl and 10 mM D,L-TBOA. All data were collected using 100 ms averaging time. Single-molecule fluorescence trajectories were selected for analysis in SPARTAN ^75^ implemented in Matlab (*Mathworks*). Trajectories were corrected for spectral bleed-through from donor to acceptor channel by subtracting a fraction of the donor intensity from the acceptor (0.168). Acquired traces were selected for analysis using the following criteria: a single catastrophic photobleaching event; over 10:1 signal-to-background noise ratio; over 5:1 signal-to-signal noise ratio; FRET lifetime of at least 5 s; a maximum of 1 donor blink per trajectory. FRET trajectories were calculated from LD555P and LD655 intensities, *I*_*D*_ and *I*_*A*_, respectively, using *E*_*FRET*_ = *I*_*A*_/(*I*_*D*_+*I*_*A*_). Population contour plots were constructed by superimposing *E*_*FRET*_ from individual trajectories and fitted to Gaussian distributions in Prism (*Graphpad*). Dwell-time distributions and transition frequencies were obtained by idealizing *E*_*FRET*_ trajectories in SPARTAN ^75^. State survival plots and dwell-time histograms were fitted to double or triple exponentials and the probability density functions ^108^, respectively. Analyses of transition frequencies, dwell times, and survival plots for individual trajectories were performed using custom made scripts implemented in Matlab (*Mathworks*). State survival plots of individual trajectories were fitted to single exponentials or double exponentials if R^2^ increased by at least 5 %, and the time constants differed at least 5-fold. Fits were only performed when traces contained at least 5 visits to the OFS or the IFS. Mean dwell times were used otherwise. For illustration purposes, state survival plots of selected molecules were refitted in Prism (*GraphPad*). 2D histograms were generated in Origin (*Originlab*). The autocorrelation coefficients *R(t/o)* of the dwell durations as functions of the number of consecutive state visits were calculated as 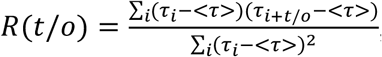, where *t/o* are the numbers turnovers, *τ*_*i*_ the dwell lengths and <*τ*> the average dwell time over the trajectory ^79^. Each experiment was performed at least in triplicate using independent protein reconstitutions and buffer preparations. In each experiment, a minimum of 400 molecules was selected.

### Transition state analysis

LFER analysis builds on observations from reaction chemistry ^30,32^, which correlate changes of the rate constants with changes of the equilibrium constant upon perturbations of the start and end equilibrium states. Here, the free energy change of the transition state, ΔG^#^ is expected to fall between the free energy changes of the equilibrium end states, the OFS and the IFS, following perturbations ^32^:

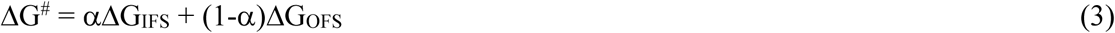

where α varies between 0 and 1. Equation 3 can be rewritten to express the energy changes relative to the OFS:

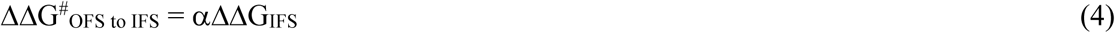

Where ΔΔG^#^_OFS to IFS_ is the activation energy of the transition from the OFS to the IFS. The activation energy of the reverse reaction can be expressed similarly:

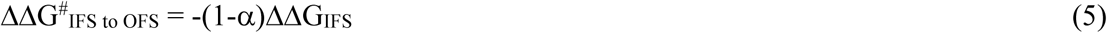

The equilibrium free energy change is calculated from equilibrium constants before and after the perturbation, *K*_*R*_ and *K*_*P*_ (where *R* and *P* stand for “reference” and “perturbation”), respectively:

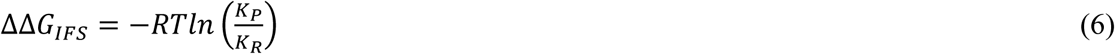

The equilibrium constants are obtained as ratios of the forward and reverse rate constants *k*_*OFS to*_ *IFS* and *k*_*IFS to OFS*_ measured before and after the perturbation. The transition state energy changes can be approximated as in Kramers theory assuming that the transition mechanism (i.e., the shape of the energy barrier) is not altered by the perturbation ^109^:

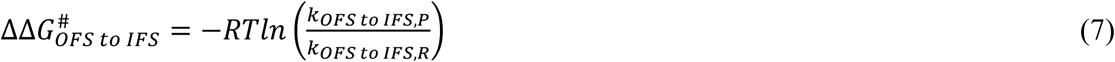

*ΔΔG*^*#*^_*IFS to OFS*_ is calculated in an analogous manner from the reverse reaction rates.

### Kinetic modeling of the transport cycle

The transport cycle for WT Glt_Ph_ was simulated in Copasi ^110^. A two-compartment system was created to reflect external and internal proteoliposome space. The initial conditions were 200 mM Na^+^ and 1 mM l-Asp in the external space with all transporters in the apo state and the fraction of the OFS of 0.6. External substrate binding and internal substrate release were approximated as irreversible. External substrate binding was arbitrarily set to 10^8^ M^-4^s^-1^, while internal release was set to an estimated value of 500 s^-1 57^. Rate constants for the conformational changes were estimated from the mean lifetimes of “typical” Y204L/K290A/A345V/V366A Glt_Ph_ molecules analyzed by smFRET and set at k_OFS to IFS, apo_ = 1.2 s^-1^, k_IFS to OFS, apo_ = 1.6 s^-1^, k_OFS to IFS, bound_ = 0.5 s^-^ 1, k_IFS to OFS, bound_ = 2.0 s^-1^. The effect of a positive allosteric modulator on k_OFS to IFS, bound_ and k_OFS_ to IFS, apo was considered the same and the increase in transport rate was simulated for 2, 5, 10, 20, 50, 100, 200, 500 and 1000 fold-increases of k_OFS to IFS, bound_ and k_OFS to IFS, apo_. Time courses were simulated over 1000 s.

## Acknowledgements

We thank Roger Altman for preparation of smFRET chambers, Zhou Zhou for dye formulations, Eva Fortea and Emma Garst for assistance with crystallization trials and Lucy Skrabanek for R scripts used in bioinformatics sequence analysis. We thank Julia Chamot-Rooke for continued support. We are grateful for the support of NINDS (R37NS085318 to O.B. and S.C.B.) and AHA (19PRE34380215 to H.D.C.). This project has received funding from the European Union’s Horizon 2020 research and innovation program under the Marie Sklodowska-Curie grant agreement MEMDYN N° 660083 (to G.H.M.H.). The structure of Y204L A345V V366A Glt_Ph_ has been submitted to the PDB (accession code 6V8G).

## Author contributions

Conceptualization: G.H.M.H. and O.B.; data acquisition: G.H.M.H.; data analysis: G.H.M.H., D.H.C., X.W., O.B.; resources: O.B. and S.C.B.; manuscript writing: G.H.M.H. and O.B. with input of all other authors

