## supplemental information for "The high-energy transition state of a membrane transporter"

#### Supplementary Tables

**Supplementary Table 1: Global difference scores (GDS) of amino acid positions with significant distribution changes.** *Related to Figure 1.*

| AA in $Glt_{Ph}$ | AA position | GDS volume | GDS physicochemical properties | AA in meso/psychrophile | Structural location |
|---|---|---|---|---|---|
| L | 13 | 0.35 | | | TM1 |
| Q | 14 | 0.78 | 0.35 | | TM1/exposed |
| I | 21 | | 0.21 | | TM1/exposed |
| V | 39 | 0.01 | | | TM1-2 loop |
| S | 65 | 0.80 | | T | TM2/TD-scaffold interface |
| A | 70 | 0.38 | | I | Intracellular hinge |
| S | 74 | 0.59 | | | Intracellular hinge |
| V | 87 | | 0.94 | | TM3/exposed |
| Y | 89 | | 0.09 | | TM3/binding |
| T | 92 | 0.06 | | | TM3/core TD |
| A | 94 | 1.41 | | T/L/I/F | TM3/HP2 packing |
| V | 97 | 1.31 | 0.55 | L/I | TM3/HP2 packing |
| R | 105 | 0.11 | | | TM3/exposed |
| I | 113 | 0.49 | | | TM3-4 loop |
| A | 127 | 0.06 | | | TM3-4 loop |
| P | 128 | 0.32 | 1.14 | | TM3-4 loop |
| P | 153 | 0.25 | 0.75 | | TM4 |
| I | 165 | 0.19 | | | TM4 |
| G | 189 | 0.19 | | | TM5/exposed |
| A | 193 | 1.05 | | | TM5/exposed |
| N | 199 | | 0.05 | | TM5/exposed |
| A | 211 | | 0.02 | | TM5 |
| V | 231 | | 0.02 | | TM6 |
| T | 232 | 0.20 | | | TM6 |
| A | 234 | 0.07 | | | TM6 |
| V | 235 | 0.84 | | F/L | TM6 |
| V | 237 | | 0.10 | | TM6/exposed |
| Q | 242 | 1.35 | 0.42 | F/L/I/V | TM6 |
| I | 243 | 0.62 | | | TM6/exposed |
| K | 254 | 0.28 | | | TM6/exposed |
| G | 255 | | 0.25 | | TM6-HP1 loop |
| P | 258 | 0.46 | | | TM6-HP1 loop |
| A | 265 | 0.48 | 0.41 | I/L/M | HP1 |
| A | 268 | 1.17 | 0.58 | E/V | HP1 |

| | | | | | |
|---|---|---|---|---|---|
| M | 269 | | 0.69 | Q/A/G/S/T | HP1 |
| T | 271 | 0.66 | 0.11 | L/I/F/V | HP1 |
| S | 279 | 0.31 | | | HP1 tip |
| T | 281 | | 0.40 | V/A | HP1 |
| V | 284 | | 0.57 | L/R/K | HP1 |
| T | 285 | 0.15 | | L/I/M/V | HP1 |
| V | 288 | 0.02 | 0.42 | K/T/A | HP1 |
| K | 289 | 1.10 | 0.51 | L/M/V | HP1 |
| M | 292 | 0.92 | 0.02 | | HP1-TM7 loop/exposed |
| A | 307 | 1.41 | 0.42 | | TM7/binding |
| Q | 318 | 0.44 | 0.37 | L/I/P | TM7/HP2 packing |
| G | 319 | 0.17 | | T/S | TM7/HP2 packing |
| V | 320 | 0.13 | | | TM7 |
| G | 330 | 0.22 | | | TM7-HP2 loop/exposed |
| T | 344 | 0.98 | | V/I | HP2 |
| A | 345 | 1.12 | 0.31 | L/S/V | HP2 |
| V | 346 | 0.38 | 0.07 | T/M/L/I | HP2 |
| A | 348 | 0.22 | 0.28 | T/S | HP2 |
| P | 356 | | 0.10 | | HP2 tip |
| V | 366 | 0.39 | | T/A | HP2 |
| S | 369 | 1.17 | | Q/V/L/M | HP2 |
| V | 370 | 0.27 | | | HP2 |
| D | 405 | | 0.57 | | TM8/binding |
| L | 406 | | 0.41 | A/S/C | TM8/HP1 packing |
| G | 408 | 0.46 | 0.50 | A/V | TM8 |
| T | 409 | 0.47 | 0.45 | | TM8 |
| T | 415 | | 0.00 | | TM8/exposed |

The table is shaded according to the structural regions as in Figure 1.

**Supplementary Table 2: Kinetic parameters of $^{3}$H-L-Asp uptake by Glt$_{Ph}$ variants.** *Related to Figure 1.*

| Glt$_{Ph}$ variant | $K_M$[a] | Fold rate change[a] |
|---|---|---|
| WT | 0.50 ± 0.04 | (average rate 0.06 ± 0.05 $s^{-1}$)[b] |
| *HP1 substitutions* | | |
| A265L | 0.42 | 1.4[c] |
| A265M | 0.34 | 1.4 |
| A265V | 0.55 | 0.6 |
| A268V | 0.30 ± 0.08 | 0.7 ± 0.5 |
| M269A | 0.59 ± 0.27 | 1.7 ± 0.5 |
| M269V | 0.77 | 1.6 |
| M269Q | 0.11 | 1.5 |
| M269L | 0.37 | 0.5 |
| T285V | 0.34 ± 0.02 | 1.8 ± 0.3 |
| A289L | 1.28 | 1.0 |
| *HP2 substitutions* | | |
| I341A | 0.27 | 0.4 |
| **A345V** | **0.79 ± 0.28** | **4.0 ± 1.2** |
| **A345S** | **0.84** | **3.8** |
| A345L | 0.70 | 0.5 |
| A345F | 0.42 | 0.3 |
| V346T | 0.25 | 0.7 |
| **M362V** | **1.11** | **7.9** |
| **V366A** | **2.82 ± 1.11** | **6.8 ± 4.1** |
| V366T | 0.53 | 1.0 |
| *Scaffold substitutions* | | |
| D48N | 0.33 | 2.0 |
| A70I | 0.31 ± 0.23 | 0.8 ± 0.3 |
| P75G | 0.29 | 1.0 |
| A191N | 0.10 | 0.5 |
| Y195F | 0.58 | 0.6 |
| **Y204L** | **0.48 ± 0.26** | **3.7 ± 1.6** |
| Y204V | 0.39 | 1.0 |
| Y204F | 0.45 | 0.7 |
| Q220A | N.D.[d] | N.D.[d] |
| H223D | 1.49 | 2.0 |
| G226L | 0.55 ± 0.72 | 1.9 ± 0.3 |
| E227G | 0.73 | 0.2 |
| *Transport domain core substitutions* | | |
| I85A | 0.25 | 1.5 |
| A94T | 0.48 | 0.7 |
| V97I | 3.01 | 3.0 |
| V97A | 0.33 | 0.9 |
| V235F | 0.50 | 1.9 |
| Q242F | 0.14 ± 0.10 | 0.4 ± 0.1 |
| L303A | 0.29 | 1.2 |
| L303V | 0.31 | 0.3 |
| Q318L | 0.38 | 1.5 |
| T322A | 0.95 | 0.8 |
| L406A | 0.15 | 0.6 |
| G408A | 0.45 | 0.4 |

| | | |
|---|---|---|
| *Combination variants* | | |
| **A345V V366A** | **1.56 ± 0.40** | **11.1 ± 5.9** |
| **Y204L A345V V366A** | **1.61 ± 0.42** | **12.7 ± 2.5** |
| D48A Y204L | 0.43 | 2.2 |
| V366A Y204L | 1.29 ± 0.69 | 5.1 ± 5.2 |
| A345V Y204L | 1.38 ± 0.60 | 3.0 ± 1.0 |
| V366A G226L | 12.55 | 5.2 |
| V366A Y204L G226L | 0.97 | 9.1 |
| A345V G226L V366A | 1.48 | 3.3 |
| V366A A345V A70I | 4.35 | 8.0 |
| A265M L406A V288T A289L | 0.62 | 1.0 |
| A345V A265M | 0.83 | 3.0 |
| Y204L A265M | 1.09 | 4.3 |
| V366A A265M | 1.57 | 2.6 |
| V366A Y204L A265M | 0.92 | 6.0 |
| V366A A345V M269A | 2.56 | 3.3 |
| V366A A345V A289L | 5.99 | 2.7 |
| V366A A345V Y204L A268V | 5.20 | 8.6 |
| V366A A345V A265M | 2.15 ± 1.54 | 11.0 ± 7.8 |
| V366A Y204L A345V M269A | 2.1 | 14.0 |
| V366A A345V Y204L A265M L406A V288T A289L | N.D.[d] | N.D.[d] |
| V366A A345V Y204L A265M | 3.1 | 3.8 |
| V366A A345V A265M G226L | 1.49 | 3.3 |
| V366A A345V A265M A70I | 2.14 | 3.3 |
| *Combinations containing K290A or R276S M395R* | | |
| R276S | 0.76 | 2.2 |
| **K290A** | **0.41 ± 0.11** | **1.7 ± 1.2** |
| **R276S M395R** | **1.33 ± 1.16** | **8.0 ± 3.2** |
| **V366A A345V Y204L K290A** | **2.64 ± 1.41** | **17.7 ± 0.9** |
| R276S M395R V366A Y204L A345V | N.D.[d] | N.D.[d] |
| R276S M395R V366A | 4.12 | 4.0 |
| **K290A Y204L** | **0.38 ± 0.17** | **2.6 ± 0.6** |
| **K290A A345V V366A** | **0.61 ± 0.15** | **9.4 ± 4.4** |
| K290A Y204L A268V A345V V366A | 5.12 | 8.2 |
| R276S Y204L A265M A345V V366A | N.D.[d] | N.D.[d] |
| R276S Y204L M269A A345V V366A | 0.88 | 3.3 |
| R276S M395R Y204L M269A A345V V366A | N.D.[d] | N.D.[d] |
| R276S M395R Y204L G226L A345V V366A | N.D.[d] | N.D.[d] |
| R276S M395R Y204L M269A L303A A345V V366A | N.D.[d] | N.D.[d] |

[a] values obtained by fitting data to Michaelis-Menten equation. When shown, errors are standard deviations for at least three independent experiments. When errors are not shown, experiments were only performed once.
[b] from 34 independent experiments.
[c] fold change of initial rate relative to the WT transporter tested on the same day.
[d] not determined because poor transport rates preclude analysis.

**Supplementary Table 3: $Glt_{Ph}$ dynamics in the presence of the non-transportable inhibitor DL-TBOA.** *Related to Figure 3.*

| $Glt_{Ph}$ variant | transition frequency ($s^{-1}$) | Number of molecules | % molecules containing 90% of measured transitions |
|---|---|---|---|
| WT | 0.03 ± 0.01 | 1287 | 8 |
| Y204L | 0.02 ± 0.01 | 1282 | 16 |
| G226L | 0.06 ± 0.02 | 1298 | 10 |
| M269A | 0.03 ± 0.01 | 1637 | 10 |
| A345V | 0.03 ± 0.02 | 1727 | 12 |
| V366A | 0.03 ± 0.01 | 1403 | 16 |
| A345V V366A | 0.03 ± 0.01 | 1678 | 10 |
| Y204L A345V V366A | 0.03 ± 0.01 | 1586 | 16 |
| K290A | 0.06 ± 0.03 | 1933 | 13 |
| K290A Y204L | 0.10 ± 0.02 | 894 | 17 |
| K290A A345V V366A | 0.02 ± 0.02 | 1067 | 13 |
| K290A Y204L A345V V366A | 0.04 ± 0.03 | 1498 | 13 |
| R276S M395R | 0.04 ± 0.01 | 742 | 12 |

**Supplementary Table 4: X-ray crystallographic data and refinement statistics for the Y204L/A345V/V366A $Glt_{Ph}$ structure deposited at the PDB.** *Related to Figure 3.*

| | |
|---|---|
| Wavelength | 0.99997 |
| Resolution range | 47.38 - 3.375 (3.496 - 3.375) |
| Space group | P 31 2 1 |
| Unit cell | 109.416 109.416 305.227 90 90 120 |
| Total reflections | 321457 (22766) |
| Unique reflections | 30612 (2964) |
| Multiplicity | 10.5 (7.7) |
| Completeness (%) | 99.54 (98.40) |
| Mean I/sigma(I) | 10.87 (1.87) |
| Wilson B-factor | 114.03 |
| R-merge | 0.1155 (1.074) |
| R-meas | 0.1212 (1.153) |
| R-pim | 0.03574 (0.4091) |
| CC1/2 | 0.993 (0.787) |
| CC* | 0.998 (0.939) |
| Reflections used in refinement | 30565 (2956) |
| Reflections used for R-free | 1400 (156) |
| R-work | 0.2324 (0.3367) |
| R-free | 0.2574 (0.3916) |
| CC(work) | 0.818 (0.835) |
| CC(free) | 0.854 (0.667) |
| Number of non-hydrogen atoms | 8679 |
| macromolecules | 8670 |
| ligands | 9 |
| Protein residues | 1179 |
| RMS (bonds) | 0.002 |
| RMS (angles) | 0.46 |
| Ramachandran favored (%) | 98.29 |
| Ramachandran allowed (%) | 1.71 |
| Ramachandran outliers (%) | 0.00 |
| Rotamer outliers (%) | 0.00 |
| Clashscore | 5.49 |
| Average B-factor | 132.82 |
| macromolecules | 132.86 |
| ligands | 96.02 |
| Number of TLS groups | 6 |

Statistics for the highest-resolution shell are shown in parentheses.

**Supplementary Table 5: Calculated population from gaussian fitting of time-averaged population.** *Related to Figure 2 and Supplementary Figures 3 and 4.*

| Apo conditions | low $E_{FRET}$ (0.4) | Intermediate $E_{FRET}$ (0.6) | High $E_{FRET}$ (0.9) |
|---|---|---|---|
| WT | 0.73 | 0.19 | 0.08 |
| M269A | 0.71 | 0.26 | 0.03 |
| G226L | 0.42 | 0.35 | 0.23 |
| Y204L | 0.71 | 0.26 | 0.23 |
| A345V | 0.61 | 0.24 | 0.15 |
| V366A | 0.64 | 0.19 | 0.17 |
| A345V V366A | 0.80 | 0.16 | 0.03 |
| Y204L A345V V366A | 0.67 | 0.22 | 0.11 |
| K290A | 0.49 | 0.49 | 0.02 |
| Y204L K290A | 0.56 | 0.38 | 0.06 |
| K290A A345V V366A | 0.62 | 0.26 | 0.12 |
| Y204L K290A A345V V366A | 0.55 | 0.41 | 0.05 |
| R276S M395R | 0.31 | 0.68 | 0.01 |
| **$Na^+$/L-Asp bound conditions** | **low $E_{FRET}$ (0.4)** | **Intermediate $E_{FRET}$ (0.6)** | **High $E_{FRET}$ (0.9)** |
| WT | 0.81 | 0.12 | 0.07 |
| M269A | 0.84 | 0.16 | 0.03 |
| G226L | 0.54 | 0.16 | 0.30 |
| Y204L | 0.77 | 0.15 | 0.08 |
| A345V | 0.65 | 0.13 | 0.22 |
| V366A | 0.63 | 0.13 | 0.23 |
| A345V V366A | 0.80 | 0.16 | 0.04 |
| Y204L A345V V366A | 0.70 | 0.20 | 0.10 |
| K290A | 0.59 | 0.27 | 0.13 |
| Y204L K290A | 0.56 | 0.26 | 0.17 |
| K290A A345V V366A | 0.63 | 0.23 | 0.14 |
| Y204L K290A A345V V366A | 0.66 | 0.26 | 0.08 |
| R276S M395R | 0.66 | 0.08 | 0.25 |

### Supplementary Figures

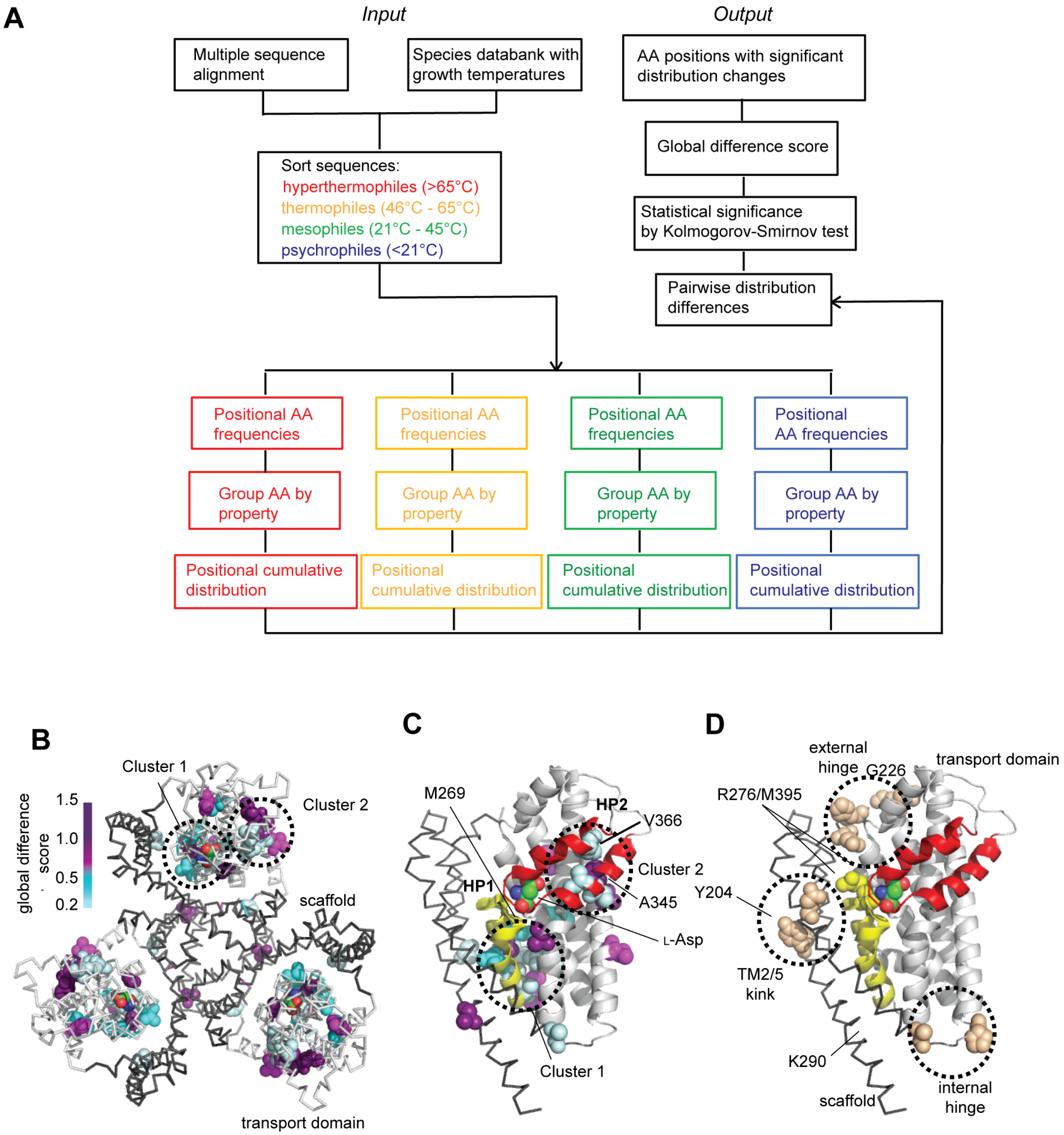

**Supplementary Figure 1: Bioinformatics analysis to identify gain-of-function mutations.** *Related to Figure 1.* (**A**) Sequence analysis workflow. (**B**) Clusters of amino acids showing changes of the sidechain volume and/or physicochemical properties correlated with the optimal growth temperature of their species of origin. The $Glt_{Ph}$ trimer is viewed from the extracellular side of the membrane and is shown in ribbon representation with the scaffold domain colored black and the transport domain colored light gray. L-Asp molecules are shown as spheres and colored according to atom type. Residues with positive global difference scores are shown as spheres and colored according to the score with the scale bar shown on the left. Dotted lines emphasize two apparent clusters in one protomer. (**C**) A single protomer viewed in the membrane plane with the transport domain shown in cartoon representation and HP1 and HP2 colored yellow and red, respectively. Substrate and candidate residues are shown as spheres and colored as in B. TM4 is removed for clarity. Dotted circles have ~10 Å diameter. (**D**) A single protomer as in C, highlighting additional mutation sites, chosen based upon structural considerations.

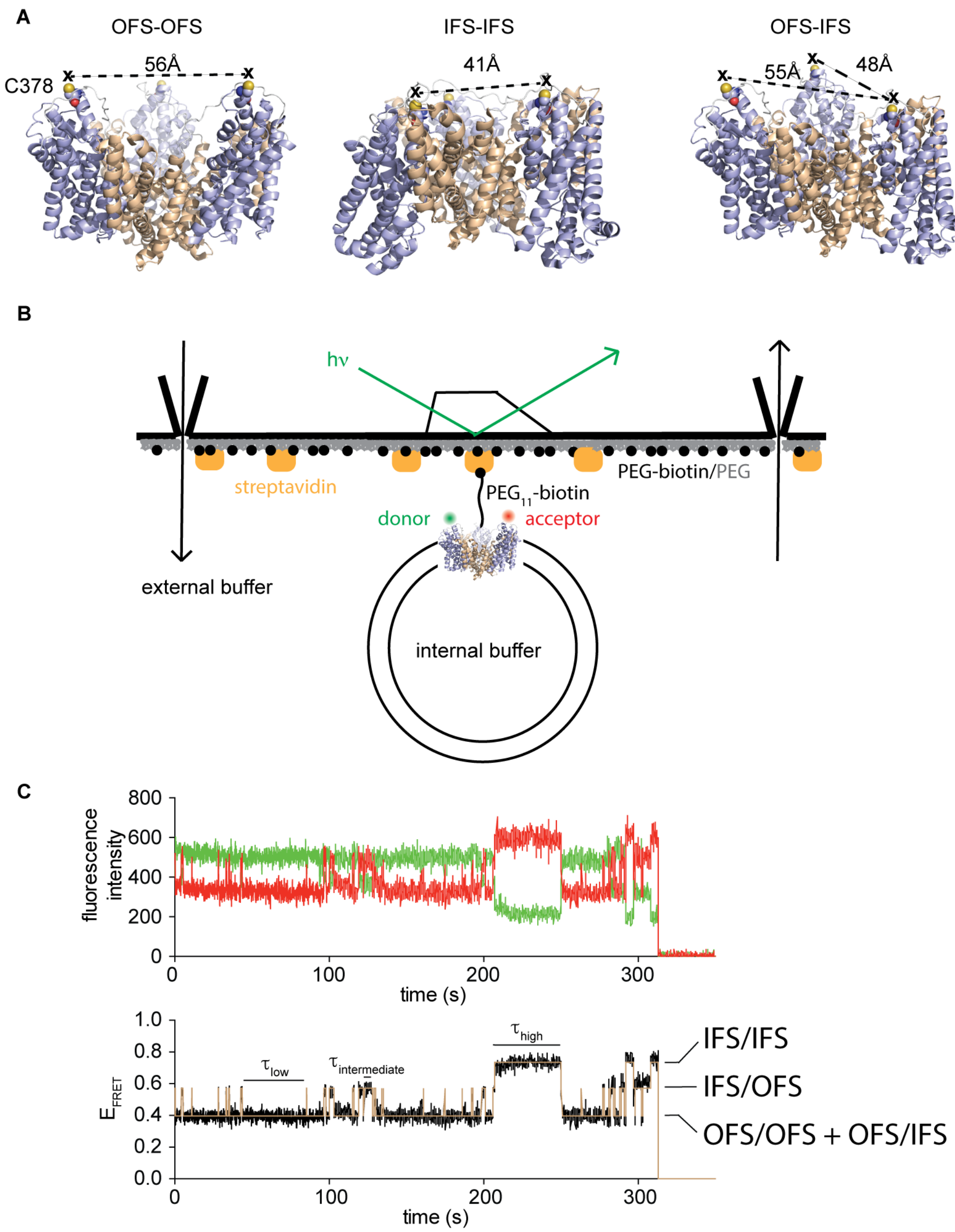

**Supplementary Figure 2: smFRET experimental setup.** *Related to Figure 2.* (**A**) Cartoon representations of the $Glt_{Ph}$ trimer labeled cysteine (by mutation of N378 to C) highlighted as spheres. Inter-subunit cysteine-to-cysteine distances are shown for two protomers in the OFS (left), in the IFS (middle) and two protomers in the OFS and the third in the IFS (right). (**B**) smFRET TIRF microscopy setup with perfusion system. (**C**) An example smFRET trajectory. Anticorrelated changes of the donor and acceptor fluorophore emissions (top) are converted into

FRET efficiencies (bottom, black) and idealized assuming three FRET efficiency ($E_{FRET}$) states (brown) corresponding to protomer configurations shown in A.

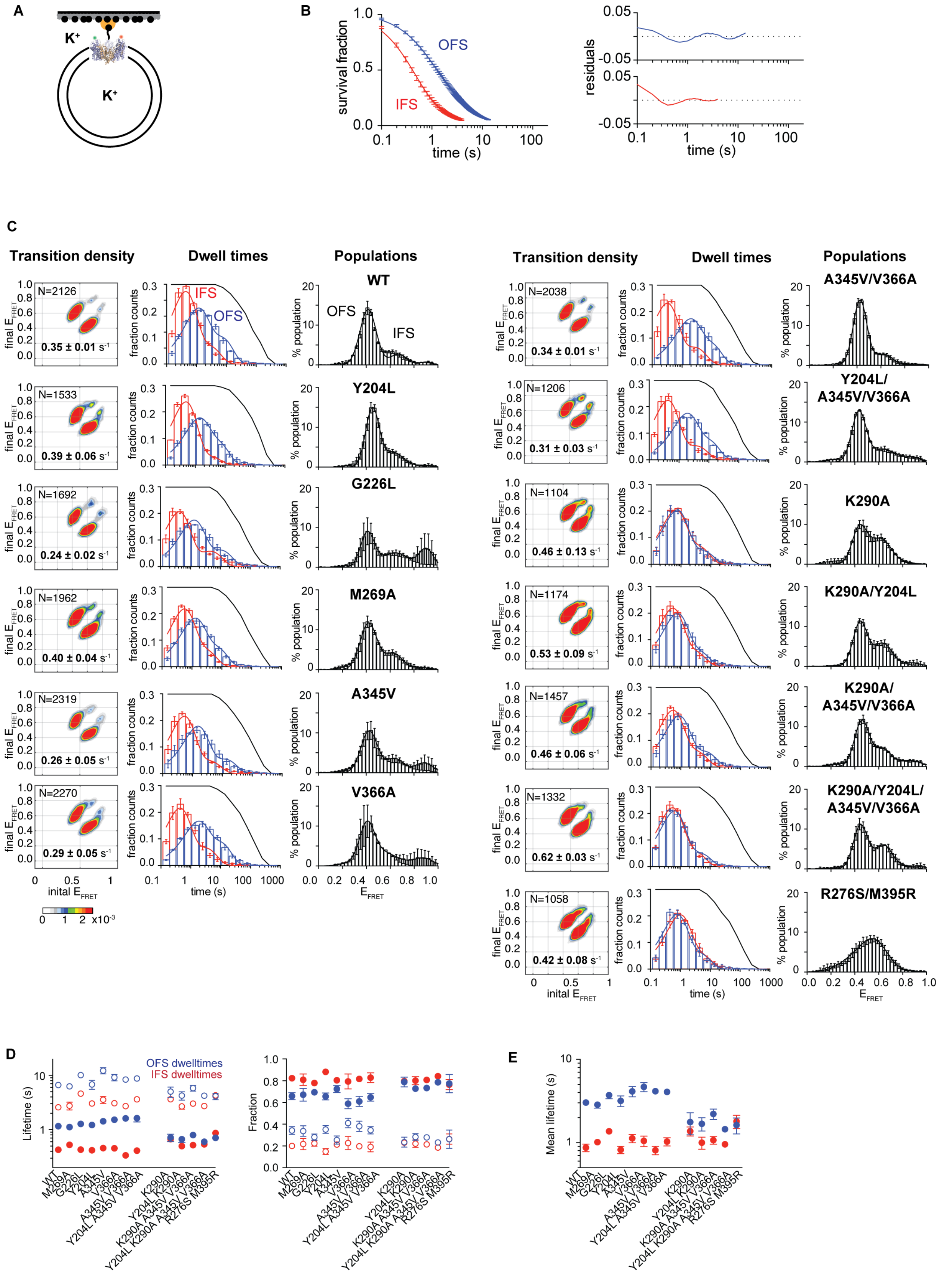

A
B
C
D
E
K+
OFS
IFS
survival fraction
time (s)
residuals
Transition density
Dwell times
Populations
WT
Y204L
G226L
M269A
A345V
V366A
A345V/V366A
Y204L/A345V/V366A
K290A
K290A/Y204L
K290A/A345V/V366A
K290A/Y204L/A345V/V366A
R276S/M395R
N=2126
0.35 ± 0.01 s-1
N=1533
0.39 ± 0.06 s-1
N=1692
0.24 ± 0.02 s-1
N=1962
0.40 ± 0.04 s-1
N=2319
0.26 ± 0.05 s-1
N=2270
0.29 ± 0.05 s-1
N=2038
0.34 ± 0.01 s-1
N=1206
0.31 ± 0.03 s-1
N=1104
0.46 ± 0.13 s-1
N=1174
0.53 ± 0.09 s-1
N=1457
0.46 ± 0.06 s-1
N=1332
0.62 ± 0.03 s-1
N=1058
0.42 ± 0.08 s-1
final EFRET
inital EFRET
fraction counts
% population
EFRET
x10-3
OFS dwelltimes
IFS dwelltimes
Lifetime (s)
Fraction
Mean lifetime (s)

**Supplementary Figure 3: Transport domain dynamics of the apo $Glt_{Ph}$.** *Related to Figure 2.* (**A**) Representation of the experimental setup. (**B**) Survival plots of the OFS and the IFS of WT $Glt_{Ph}$ fitted to double exponentials with residuals shown on the right. (**C**) For each $Glt_{Ph}$ variant, shown are the following plots. Left panels: transition density plots. Shown on the panels are the number of trajectories analyzed (*N*) and the population-wide mean transition frequency calculated as the ratio of the total number of transitions and the total trajectory lifetime. Scalebar is shown at the bottom. Middle panels: dwell-time distributions of the OFS in blue and the IFS in red. Thin lines through the data are fits to biexponential probability density functions. Black lines are the survival of the FRET lifetimes normalized between 1 and 0. Right panels: time-averaged population distributions of $E_{FRET}$ fitted to three Gaussian distributions with mean $E_{FRET}$ values of ~0.4, 0.6 and 0.9; (**D**) Lifetimes (left) and component fractions (right) for the OFS (blue) and the IFS (red) obtained by fitting dwell-time distributions in C. Open and filled symbols represent the long and short lifetime components, respectively. (**E**) Mean OFS and IFS lifetimes calculated using fitted parameters from D. Data are averages and standard errors of at least three independent measurements.

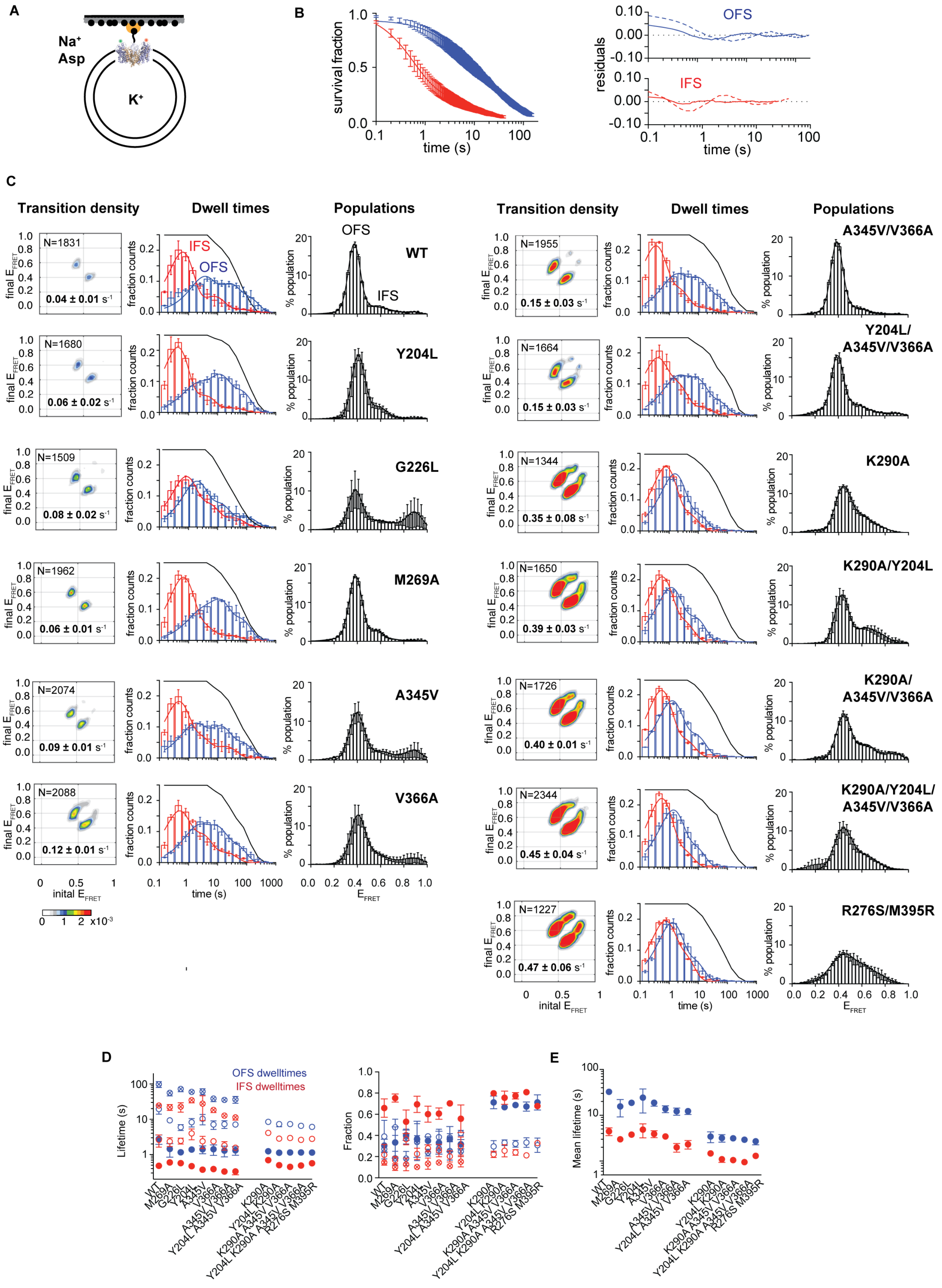
A
Na+
Asp
K+
B
survival fraction
time (s)
residuals
OFS
IFS
C
Transition density
Dwell times
Populations
final E_FRET
initial E_FRET
fraction counts
% population
E_FRET
WT
Y204L
G226L
M269A
A345V
V366A
A345V/V366A
Y204L/A345V/V366A
K290A
K290A/Y204L
K290A/A345V/V366A
K290A/Y204L/A345V/V366A
R276S/M395R
N=1831
0.04 ± 0.01 s-1
N=1680
0.06 ± 0.02 s-1
N=1509
0.08 ± 0.02 s-1
N=1962
0.06 ± 0.01 s-1
N=2074
0.09 ± 0.01 s-1
N=2088
0.12 ± 0.01 s-1
N=1955
0.15 ± 0.03 s-1
N=1664
0.15 ± 0.03 s-1
N=1344
0.35 ± 0.08 s-1
N=1650
0.39 ± 0.03 s-1
N=1726
0.40 ± 0.01 s-1
N=2344
0.45 ± 0.04 s-1
N=1227
0.47 ± 0.06 s-1
x10-3
D
OFS dwelltimes
IFS dwelltimes
Lifetime (s)
Fraction
E
Mean lifetime (s)

**Supplementary Figure 4: Transport domain dynamic under transport conditions**. *Related to Figure 2.* (**A-E**) Panels are as in Supplementary Figure 3. In B, data are fitted to triple exponentials with residuals shown on the right. Dashed lines correspond to residuals of the double exponential fits. In C, dwell-time distributions were fitted to tri-exponential probability density functions for mutants that do not contain K290A or R276S/M395R mutations. The dwell-time distributions of the latter were fitted to biexponential probability density functions. In D, crossed, open and solid symbols correspond to long, intermediate and short lifetimes, respectively.

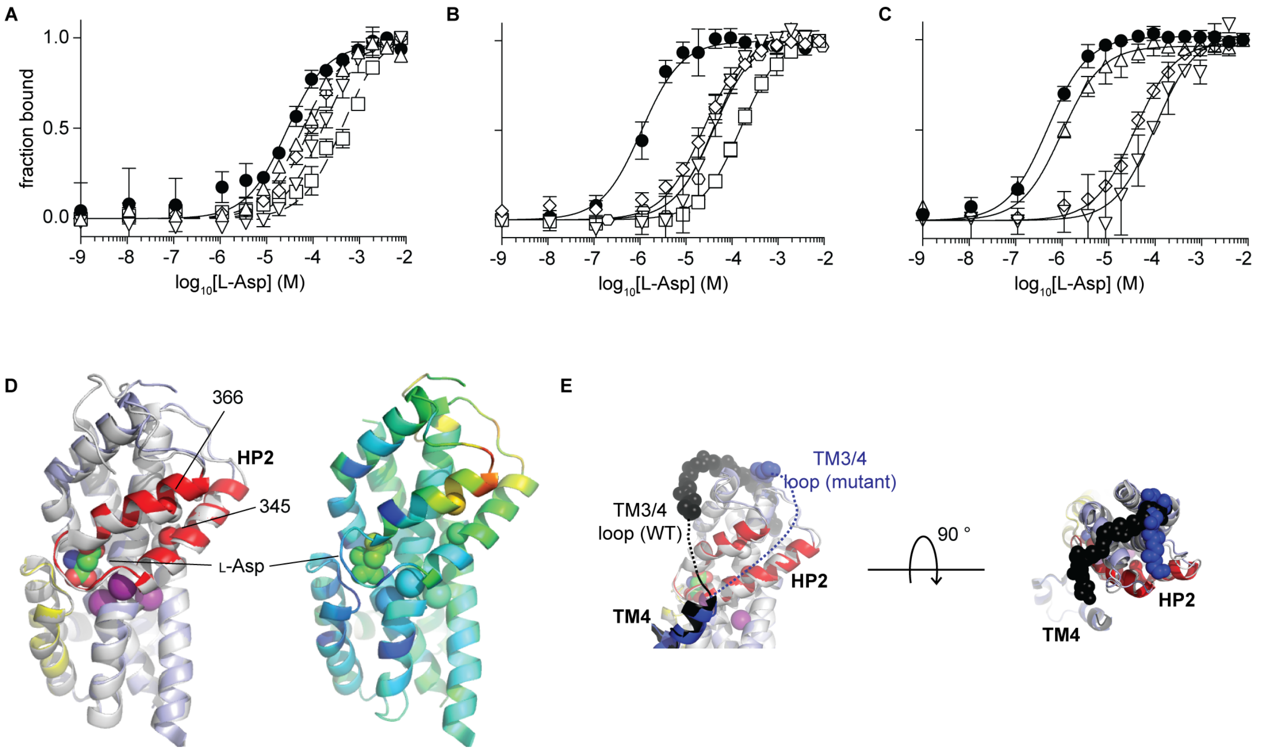

**Supplementary Figure 5: L-Asp affinity of the Glt$_{Ph}$ variants.** *Related to Figure 3.* (**A-C**) L-Asp binding isotherms measured at 25 °C in the presence of 1 mM $Na^+$ (**A**) or 10 mM $Na^+$ (**B** and **C**). Lines represent fits to the Hill equation with $n = 1$. In A: WT (filled circles), Y204L (triangles), G226L (diamonds), M269A (downward triangles), A345V (squares). In B: WT (filled circles), V366A (squares), A345V/V366A (triangles), Y204L/A345V/V366A (diamonds), R276S/M395R (hexagons). In C: K290A (filled circles), Y204L/K290A (triangles), K290A/A345V/V366A (downward triangles), Y204L/K290A/A345V/V366A (diamonds). (**D**) Superimposition of the transport domains of the Y204L/A345V/V366A mutant and WT Glt$_{Ph}$ (PDB accession code: 2NWX). The structures are colored as in Figure 3 (left) and Y204L/A345V/V366A Glt$_{Ph}$ from low (blue) to high (orange) B-factors (right). Location of residues 345 and 366 are highlighted by depicting their Cα atoms as spheres. (**E**) Close-up of the resolved N-terminal section of the TM3/4-loop by superimposition of the transport domains of Y204L/A345V/V366A mutant and the WT Glt$_{Ph}$ (PDB accession code: 2NWX).

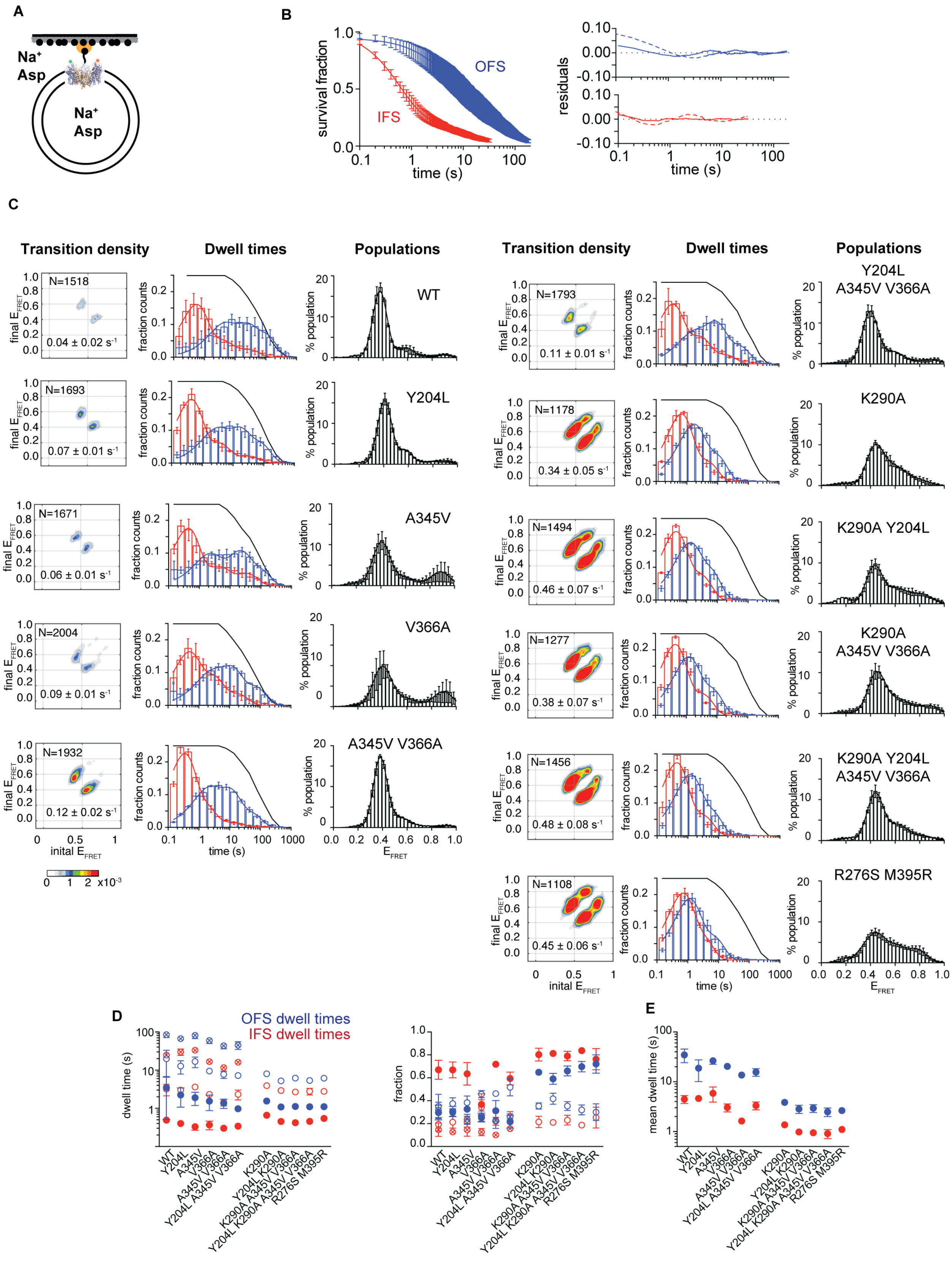

A
B
C
D
E
OFS
IFS
survival fraction
residuals
time (s)
Transition density
Dwell times
Populations
WT
Y204L
A345V
V366A
A345V V366A
Y204L A345V V366A
K290A
K290A Y204L
K290A A345V V366A
K290A Y204L A345V V366A
R276S M395R
N=1518
0.04 ± 0.02 s-1
N=1693
0.07 ± 0.01 s-1
N=1671
0.06 ± 0.01 s-1
N=2004
0.09 ± 0.01 s-1
N=1932
0.12 ± 0.02 s-1
N=1793
0.11 ± 0.01 s-1
N=1178
0.34 ± 0.05 s-1
N=1494
0.46 ± 0.07 s-1
N=1277
0.38 ± 0.07 s-1
N=1456
0.48 ± 0.08 s-1
N=1108
0.45 ± 0.06 s-1
final EFRET
inital EFRET
fraction counts
% population
EFRET
OFS dwell times
IFS dwell times
dwell time (s)
fraction
mean dwell time (s)

**Supplementary Figure 6: Transport domain dynamics under symmetric conditions in the presence of 200 mM $Na^+$ ions and 100 µM L-Asp**. *Related to Figure 4 and 5.* (**A-E**) Panels are as in Supplementary Figure 4.

Figure S7

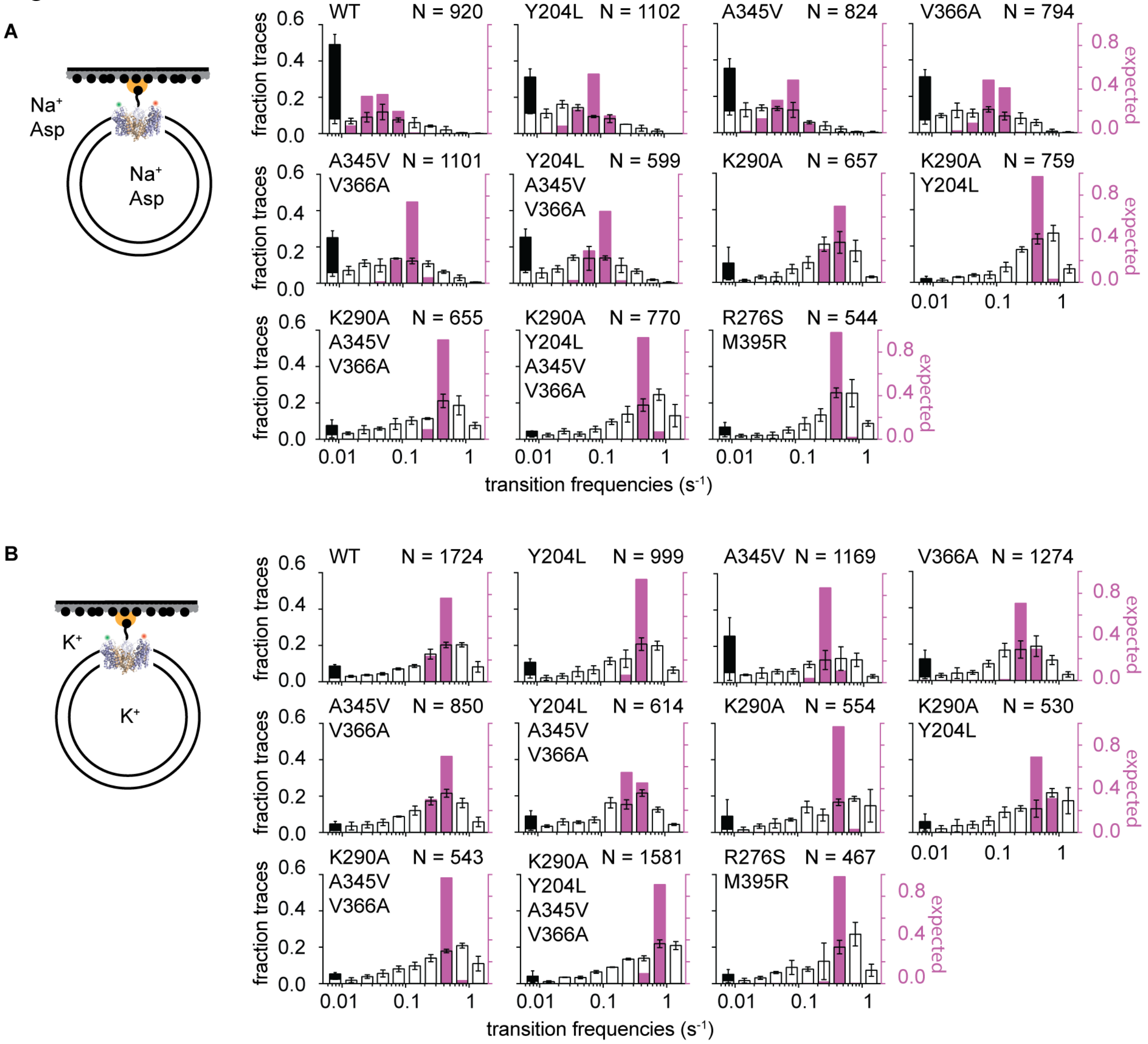

**Supplementary Figure 7: Distributions of transition frequencies of individual trajectories in the presence of symmetric 200 mM Na$^+$ ions and 100 µM L-asp (A) and under apo conditions (B).** *Related to Figure 4.* The open bars show the transition frequency distribution of Glt$_{Ph}$ molecules with at least one transition. The stacked black bars show the fractions of the trajectories without transitions. The pink bars show the expected binomial distributions calculated assuming that all trajectories share the same mean transition frequency as reported in Supplementary Figure 6. The number of trajectories included in the analysis, *N* is shown on the panels. Data are shown as averages and standard errors of three independent experiments.

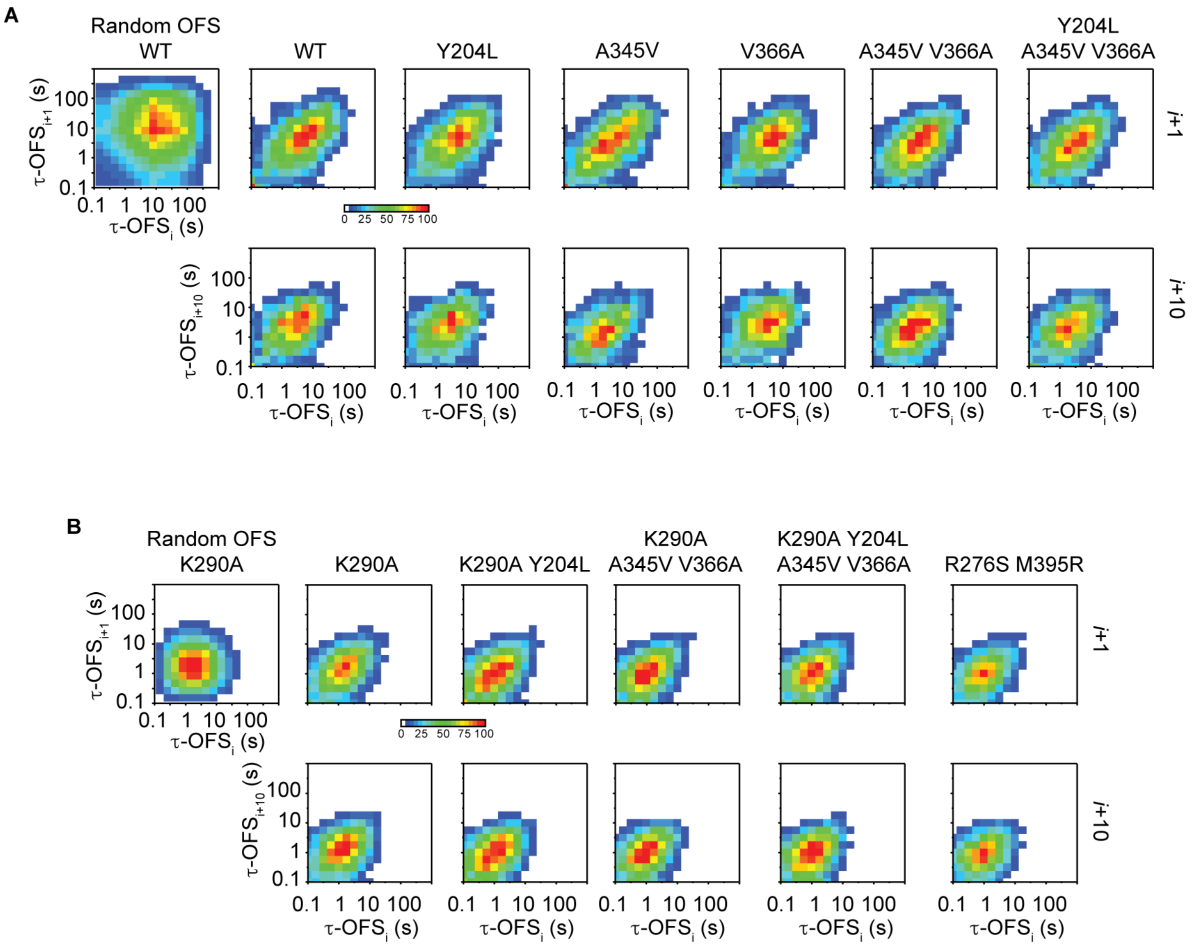

**Supplementary Figure 8: Correlated durations of sequential OFS dwells in $Glt_{Ph}$ variants.** *Related to Figure 4.* 2D histograms of durations of two consecutive OFS dwells (top row) and two OFS dwells separated by 10 visitations to the state (bottom row) are shown for less (**A**) and more (**B**) dynamic $Glt_{Ph}$ variants. Histograms expected for random uncorrelated dwells are shown on the left for the WT $Glt_{Ph}$ (**A**) and the K290A mutant (**B**). Characteristic elongated shapes of the experimental histograms are due to the correlation between the lengths of the dwells.

**A**

| $Glt_{Ph}$ variant | % traces OFS | % traces IFS |
|---|---|---|
| WT | 95 ± 1 | 97 ± 1 |
| Y204L | 88 ± 2 | 94 ± 1 |
| A345V | 90 ± 5 | 95 ± 2 |
| V366A | 89 ± 1 | 94 ± 2 |
| A345V V366A | 89 ± 2 | 92 ± 1 |
| Y204L A345V V366A | 83 ± 1 | 89 ± 1 |
| K290A | 82 ± 7 | 87 ± 2 |
| K290A Y204L | 86 ± 3 | 88 ± 1 |
| K290A A345V V366A | 86 ± 2 | 88 ± 2 |
| K290A Y204L A345V V366A | 86 ± 1 | 88 ± 2 |
| R276S M395R | 81 ± 2 | 88 ± 1 |

**B**

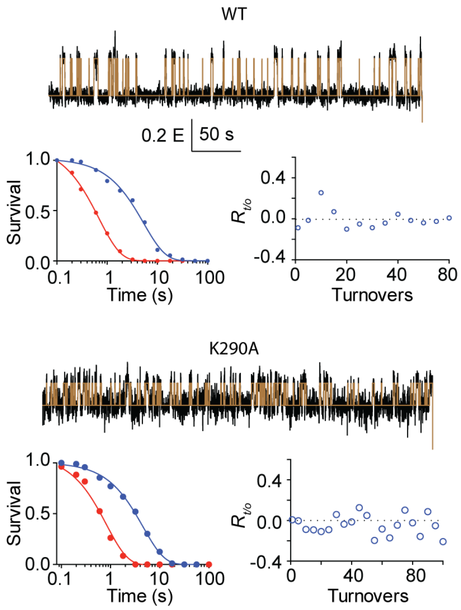

**C**

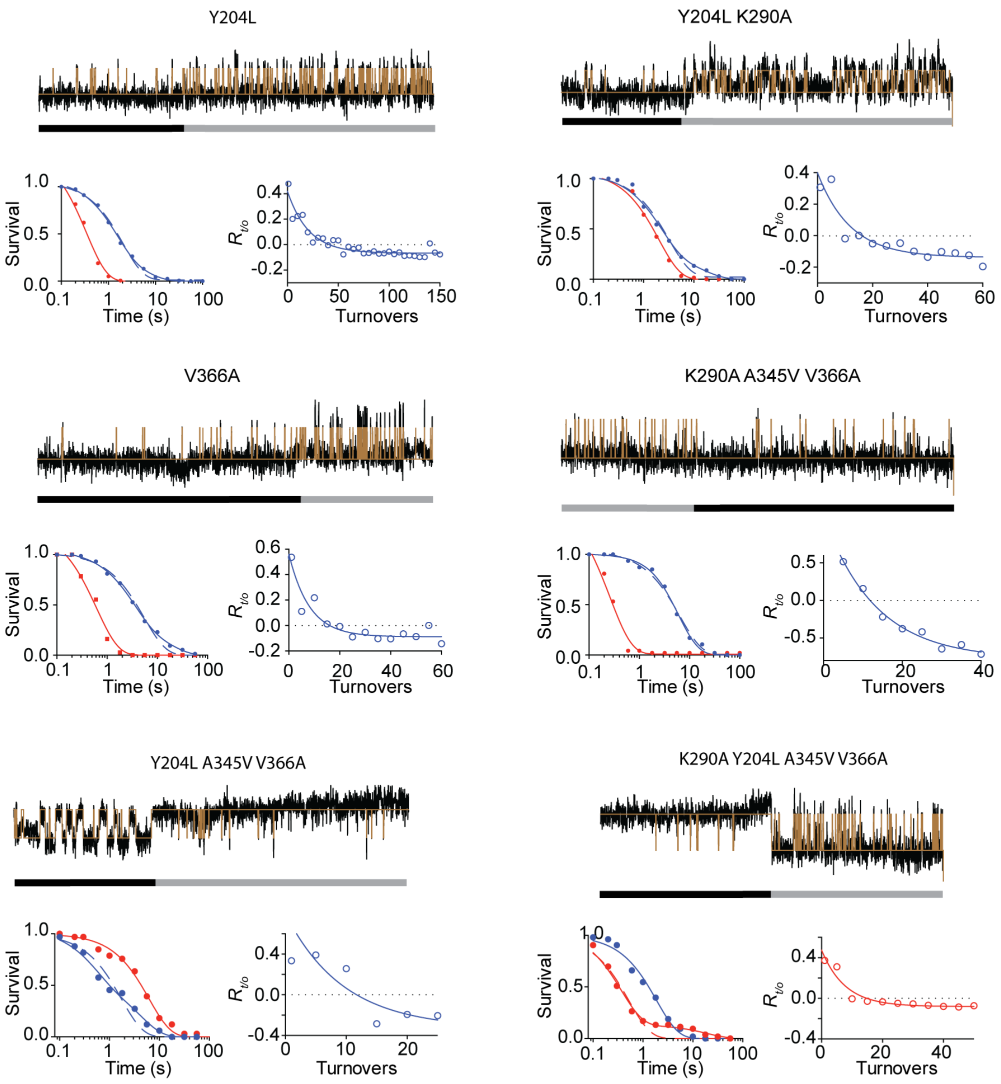

**Supplementary Figure 9: Mode switching.** *Related to Figure 4.* (**A**) Fractions of kinetically homogeneous traces for which the survival of OFS and IFS dwell times fit well to single exponential functions. (**B**) Examples of kinetically homogeneous single-molecule trajectories recorded for the WT $Glt_{Ph}$ and the K290A mutant in the presence of 200 mM $Na^+$ ions and 100 μM L-Asp. Below each trajectory, shown are the survival plots for the OFS (blue) and the IFS (red) fitted to single exponential functions and the plots of autocorrelation coefficients, $R_{t/o}$, fluctuating around 0. (**C**) Traces showing mode switching. Black and grey bars below the trajectories indicate apparent slow and fast modes. For these traces, fitting the survival plots of either the OFS, IFS, or both to double exponential functions (solid lines) improves the coefficient of determination over 5 % compared to single exponentials (dotted lines). Autocorrelation functions reflect temporal segregation of the similar dwells. Solid lines are fits to single exponentials to guide the eye.

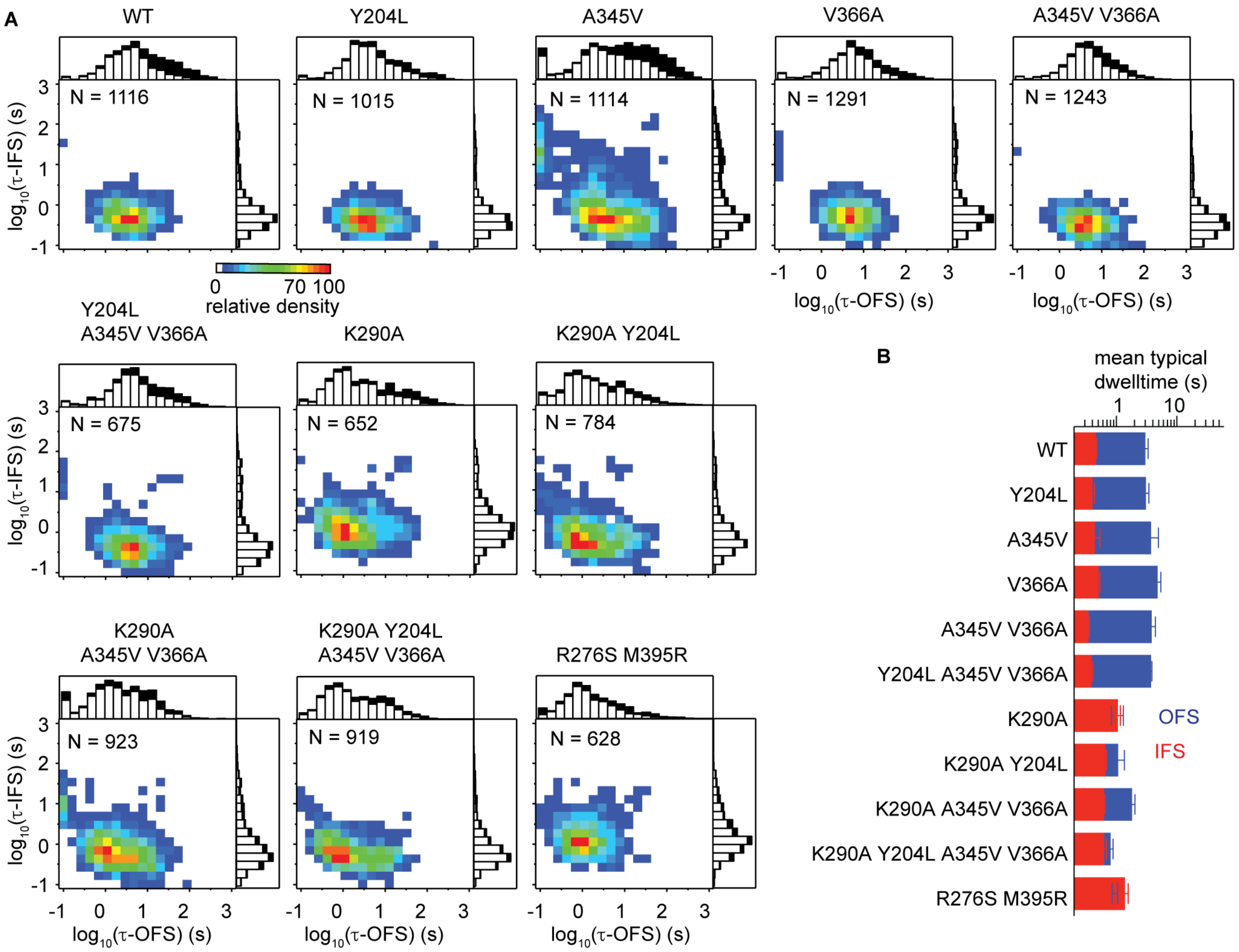

**Supplementary Figure 10: Kinetic heterogeneity of Glt$_{Ph}$ variants under apo conditions.** *Related to Figure 6.* **(A)** 2D histograms of the OFS and the IFS lifetime pairs obtained for all trajectories exhibiting single exponential behavior of the survival plots. The number of traces *N* is shown on the panels. Stacked histograms of the mean lifetimes (open bars) and the molecules showing no transitions before photobleaching (black bars) are shown above (of the OFS) and to the right (for the IFS) of the panels. Scale bar shows relative density normalized to the number of molecules included (*N*). (**B**) Mean lifetimes of the molecules falling within 30 % of the most populated bins of the 2D-histograms (yellow to red). Data are shown as means and standard errors of three independent measurements. All data were combined to construct the 2D histograms in A for presentation purposes.

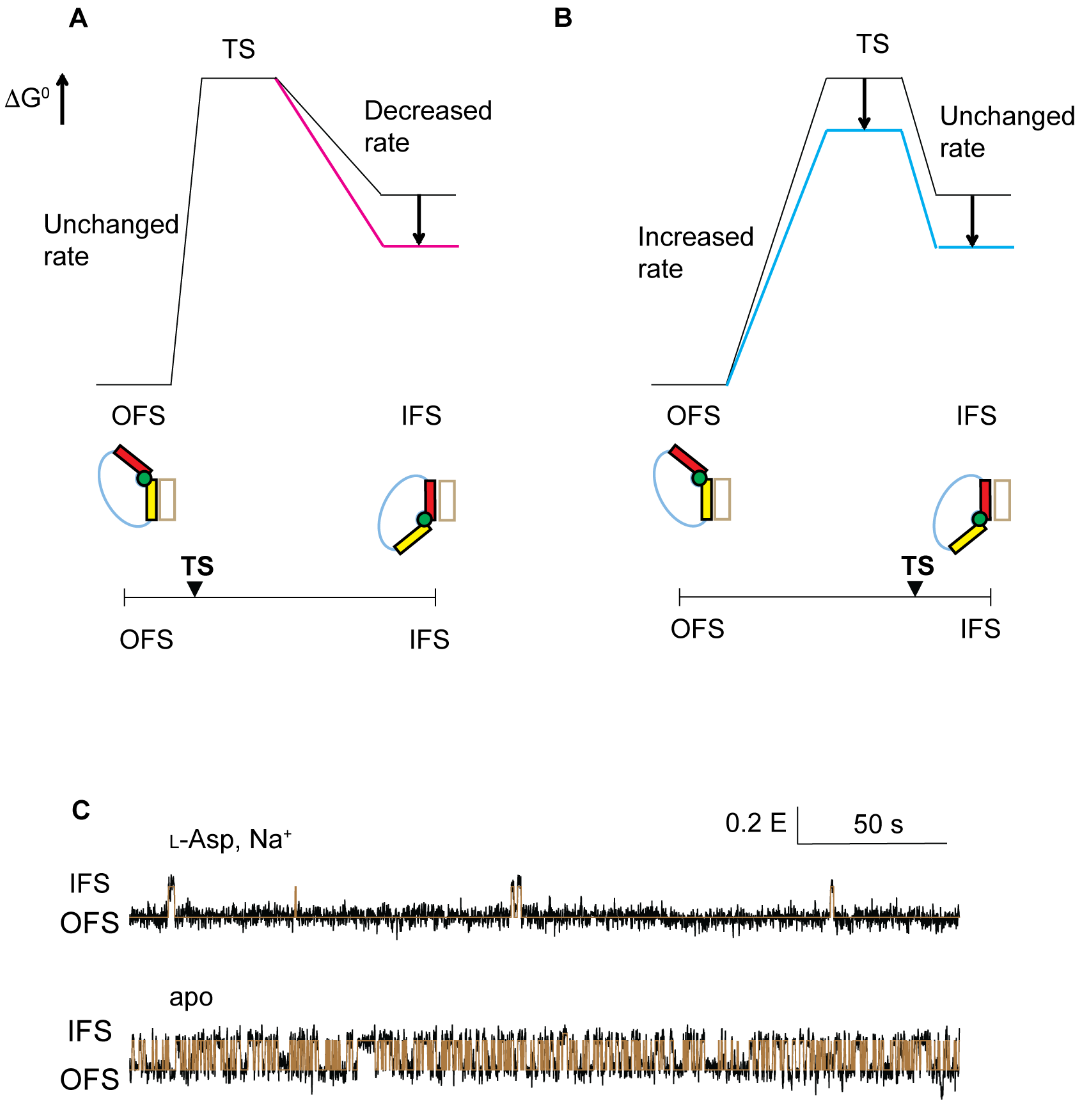

**Supplementary Figure 11: Transition state analysis.** *Related to Figure 6.* (**A, B**) Free energy diagram of the OFS to IFS conformation change (black) and the expected effect of perturbations (colors) if the TS is similar to the OFS (**A**) or the IFS (**B**). Cartoon representations of the low-energy states are below the diagrams. (**C**) Representative smFRET trajectories for WT $Glt_{Ph}$ bound to L-Asp and $Na^+$-ions (top) and in the absence of ligands (bottom). Scale bar is on the upper right of the panel.
